# Mechanisms of transforming DNA uptake to the periplasm of *Bacillus subtilis*

**DOI:** 10.1101/2021.03.09.434615

**Authors:** Jeanette Hahn, Micaela DeSantis, David Dubnau

**Author notes:** Public Health Research Institute and Department of Microbiology, Biochemistry and Molecular Genetics, New Jersey Medical School, Rutgers University (David Dubnau). Address correspondence to David Dubnau.

## Abstract

We demonstrate here that the acquisition of DNAase resistance by transforming DNA, often assumed to indicate transport to the cytoplasm, actually reflects uptake to the periplasm, requiring a re-evaluation of conclusions about the roles of several proteins in transformation. The new evidence suggests that the transformation pilus is needed for DNA binding to the cell surface near the cell poles and for the initiation of uptake. The cellular distribution of the membrane-anchored ComEA of *B. subtilis* does not noticeably change during DNA uptake as does the unanchored ComEA of *Vibrio* and *Neisseria*. Instead, our evidence suggests that ComEA stabilizes the attachment of transforming DNA at localized regions in the periplasm and then mediates uptake, probably by a Brownian ratchet mechanism. Following that, the DNA is transferred to periplasmic portions of the channel protein ComEC, which plays a previously unsuspected role in uptake to the periplasm. We show that the transformation endonuclease NucA also facilitates uptake to the periplasm and that the previously demonstrated role of ComFA in the acquisition of DNAase resistance actually derives from the instability of ComGA when ComFA is deleted. These results prompt a new understanding of the early stages of DNA uptake for transformation.

**IMPORTANCE:** Transformation is a widely distributed mechanism of bacterial horizontal gene transfer that plays a role in the spread of antibiotic resistance and virulence genes and more generally in evolution. Although transformation was discovered nearly a century ago and most, if not all of the proteins required have been identified in several bacterial species, much remains poorly understood about the molecular mechanism of DNA uptake. This study uses epifluorescence microscopy to investigate the passage of labeled DNA into the compartment between the cell wall and the cell membrane of *Bacillus subtilis*, a necessary early step in transformation. The roles of individual proteins in this process are identified, and their modes of action are clarified.

## INTRODUCTION

Transformation, the ability to internalize high molecular weight environmental DNA, is widespread among bacteria and is a major mechanism of horizontal gene transfer (reviewed in (1–3)). Several proteins that mediate transformation are widely conserved, suggesting that the core mechanism is of ancient origin. Although most of these proteins were identified more than three decades ago in a genetic screen (4, 5), much remains to be learned of the mechanisms that mediate the transfer of DNA across the multiple layers that comprise the bacterial surface barrier.

In the Gram-negative bacteria *Vibrio cholerae* and *Neisseria gonorrhoeae*, a type 4 pilus (t4 pilus) snares transforming DNA (tDNA) and retracts, threading the tDNA through a pore in the outer membrane formed by a ring of secretin subunits and thus into the periplasm (6–9). There, the tDNA encounters the DNA-binding protein ComEA, a periplasmic protein that serves as a Brownian ratchet (10), preventing back diffusion of the tDNA which consequently is believed to accumulate in the periplasm before crossing the inner cell membrane (7–9, 11, 12). In these bacteria, ComEA diffuses to the site of transforming tDNA uptake where it concentrates dramatically as it binds to inward-diffusing tDNA.

Gram-positive bacteria lack an outer membrane. Instead, *Bacillus subtilis*, the subject of the present study, is surrounded by a formidable cell wall, roughly 30-35 nm in thickness, enclosing a 20-40 nm thick periplasm, defined as the compartment between the wall and the cell membrane (13–15). This compartment is probably gel-like, containing proteins, small molecules and membrane-anchored lipoteichoic acid (14, 16), all of which may restrict diffusion by molecular crowding. The wall of *B. subtilis* is a complex structure, including peptidoglycan, proteins and wall teichoic acid (14). These differences in the outer layers of Gram-positive and Gram-negative bacteria suggest consequent differences in their mechanisms of DNA uptake to the periplasm. In fact, wall teichoic acid, which is exclusive to the Gram positives, has been proposed to play a role in DNA binding to competent cells (17). Also suggestive of such differences is that *B. subtilis* ComEA is an integral membrane protein with its C-terminal DNA binding motifs exposed to the periplasm (18, 19) as opposed to the unanchored ComEA proteins of *V. cholerae* and *N. gonorrhoeae* that can diffuse within the periplasm.

Transformable Gram-positive bacteria, like their Gram-negative counterparts, encode t4pilus-like proteins that form filamentous structures. In *Streptococcus pneumoniae*, these proteins assemble fibers that can extend several micrometers into the extracellular environment and can bind DNA (20–22). No such extended structure has been described in *B. subtilis*, which instead seems to form a shorter “pseudopilus” that is probably long enough to traverse the wall (23). This transformation pseudopilus (tpilus) is encoded by the 7 genes of the *comG* operon and also requires *comC* and *bdbDC* for its construction (23–28). ComC is a membrane peptidase that processes ComGC, the major pre-pilin subunit as well as several minor pre-pilins, while BdbD and BdbC are thiol-disulfide oxidoreductases that introduce intramolecular and intermolecular disulfide bonds into ComGC and ComEC, the latter a component of the transformation permease (29). *comGA*, the first gene of the *comG* operon, encodes a traffic ATPase that is required for tpilus assembly (30). By analogy with *V. cholerae* (6) it is likely that the Gram-positive tpili retract to bring DNA into the periplasmic compartment. Indeed, a recent study shows convincingly that the pili of *S. pneumoniae* extend and rapidly retract (22), although no such evidence has been reported for *B. subtilis*.

In all transformable bacteria, transport across the cell membrane is accomplished with the participation of ComEC, which is believed to form a channel for the passage of DNA. In *B. subtilis* and *S. pneumoniae*, transport also requires the membrane associated ComFA ATPase and ComFC, its binding partner (31–35). During transport, one strand of the transforming tDNA is degraded while the transforming strand enters the cytosol (36), where it associates with DNA binding proteins and recombines with homologous DNA sequences to yield a transformant.

In both Gram-negative and Gram-positive bacteria, tDNA becomes resistant to added DNAase I (hereafter DNAase) during an early step in transformation. In the Gram-negatives, DNAase-resistance signals that the tDNA has breached the outer membrane. In Gram-positive bacteria the proper interpretation of DNAase resistance is less clear and has been equated with “uptake” or “internalization” (37–39) without precise definition, although often assumed to indicate transport to the cytoplasm. The latter assumption was reasonably suggested by the observations that ComFA and ComEC, both required for the acquisition of DNAase resistance, are almost certainly needed for transport to the cytoplasm, based on their membrane localizations and molecular properties.

This study elucidates the process of tDNA uptake into the periplasm of *B. subtilis*. We show here that DNAase resistance actually reflects entry of tDNA to the periplasm, a step that we refer to as “uptake” to distinguish it from “transport” across the cell membrane. With this understanding, we have visualized the process of uptake using fluorescently labeled tDNA, enabling a re-evaluation of the roles of several transformation proteins. Based on our results we present an updated understanding of the early steps in transformation of *B. subtilis*, which may be applicable to other transformable firmicutes.

## RESULTS

### Experimental design

To visualize uptake, bacteriophage lambda DNA (48.5 kbp) was labeled using the Label-IT Rhodamine TM reagent (Mirus Bio) that covalently couples the fluorophore predominantly to the N7-position of guanine through a flexible linker, without disrupting Watson-Crick base pairing. To minimize perturbation of the rhodamine-DNA (rDNA) structure we used a labeling density that modifies fewer than 1% of the bases.

In *B. subtilis*, competence for transformation is expressed in only 10-20% of the cells. To identify competent cells, the promoter of *comG* was fused at the *amyE* locus to either yellow fluorescent protein (YFP) or cyan fluorescent protein (CFP), which delineate the cytosolic compartments of the competent cells. These isogenic strains behaved identically in our assays. For each experiment, aliquots of YFP- and CFP-expressing competent cells were incubated separately with rDNA, fixed with paraformaldehyde to stop transformation and then combined and visualized for epifluorescence and phase contrast microscopy. Depending on the experiment, the YFP- and CFP-expressing samples were derived from wild-type and mutant strains or from samples incubated with and without DNAase after transformation. This permitted direct comparisons without differences due to image processing and the vagaries of agarose pads.

### DNAase as a tool for transformation studies

Studies in *B. subtilis* and *S. pneumoniae* have used 50-200 μg/ml of DNAase under varying conditions to investigate transformation (4, 19, 34, 37, 38, 40). Because we wished to study uptake, we sought to standardize a DNAase treatment that would remove attached rDNA from the surface of the cells, without degrading rDNA in the unidentified protected compartment. Preliminary experiments comparing treatment with DNAase concentrations from 10 to 100 μg/ml after 15 minutes of incubation with rDNA, when the acquisition of DNAase resistance has reached about half its maximum level (37), showed no differences in the intensities of the rhodamine signals after DNAase treatment, comparing the lowest and highest amounts of DNAase. If DNAase significantly penetrated the protected compartment, we would have expected to see a dose-dependent decrease in the residual rDNA signal. We therefore settled on a 3-minute incubation with 100 μg/ml of DNAase at 37°C as an appropriate method for our experiments.

Wild-type cultures expressing YFP or CFP were each incubated with 0.2 μg/ml rDNA, close to a saturation concentration for transformation. The CFP samples were treated with DNAase and were then mixed with untreated YFP cells from the same time points and visualized for fluorescence in the YFP, CFP and rhodamine channels as well as by phase contrast after 2- and 30-minutes incubation with rDNA. In the 2-minute sample (Figure 1A), few competent cells and very few of the non-competent cells were associated with rDNA and after DNAase treatment virtually no cells retained a significant rhodamine signal. After 30-minutes incubation (Figure 1B), most of the competent cells showed an rDNA signal without DNAase treatment while the non-competent cells were rarely (<1%) associated with rDNA, showing that association is biologically relevant. After DNAase treatment, many of the cells from the 30-minute sample were still associated with rDNA (Figure 1B). The data presented to this point, are consistent with a process in which DNA first binds specifically to the surface of competent cells in a DNAase sensitive fashion and later becomes DNAase resistant by entering a protected compartment. DNAase resistance is largely complete at 30 minutes, consistent with previous observations using radiolabeled tDNA (37), suggesting that the uptake of rDNA is not markedly impeded by its covalent adducts.

**Figure 1.**
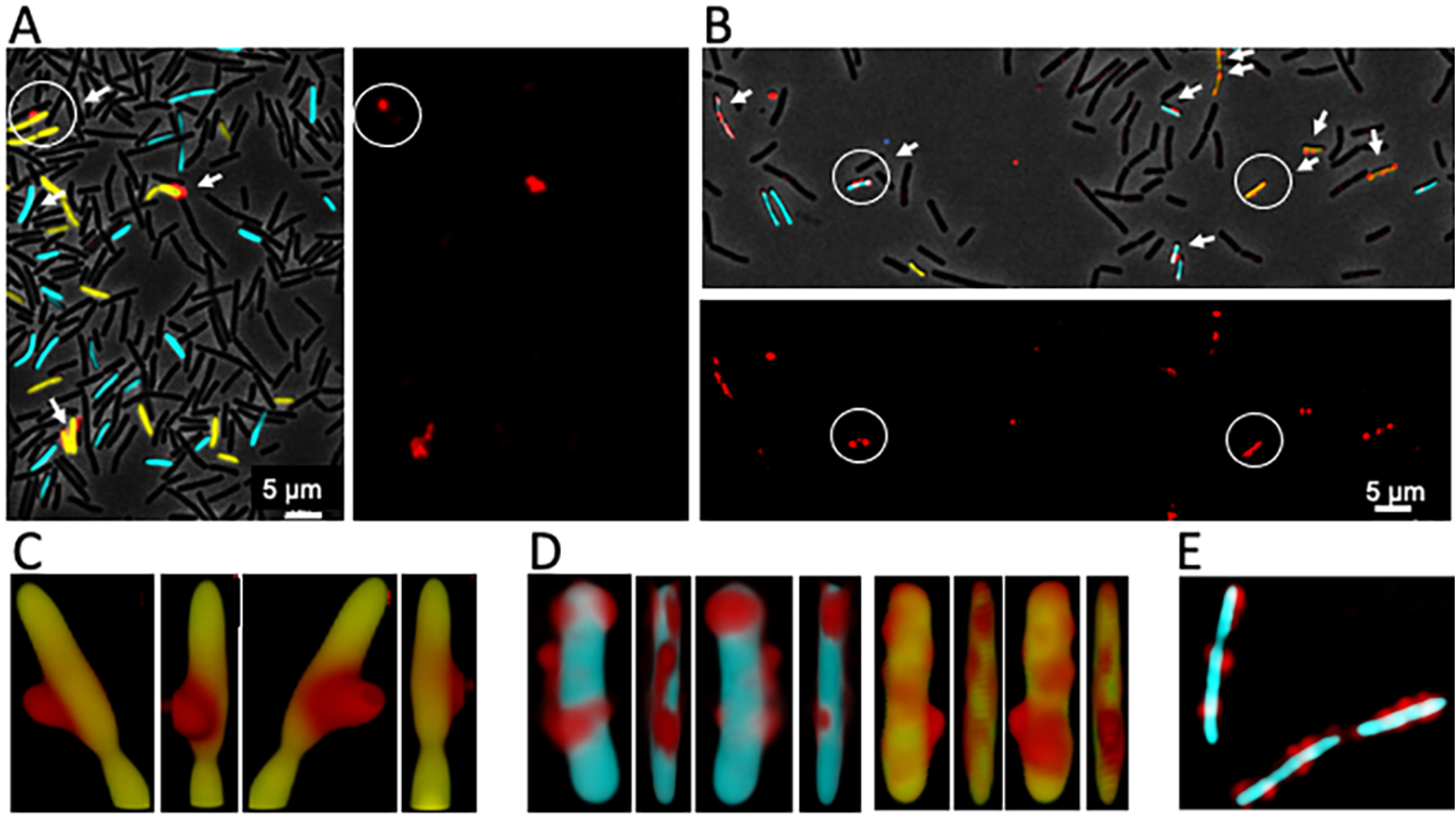
Binding of tDNA and acquisition of DNAase resistance. Competent cells expressing CFP (BD5810) or YFP (BD6011) from the *comG* promoter were incubated separately with rDNA and samples were taken after (A) 2- and (B) 30-minutes of incubation. The CFP samples were treated with DNAase and then combined with YFP samples from the same time points. In panels A and B, images are presented with superimposed rhodamine, CFP, YFP and phase contrast channels as well as one showing only the rhodamine channel for clarity. Cells associated with rDNA are indicated by arrows. The circled cell in panel A was imaged by 3D deconvolution and volume reconstructed and views produced by successive 90° rotations are shown in panel C. In panel D volume reconstructed images are shown for the single cyan (with DNAase) and yellow (no DNAase) cells, circled in panel B. Panel E contains a 3D-deconvolved optical slice from the center of a Z-stack, showing of a group of cells after DNAase treatment.

### rDNA enters the periplasm where it becomes resistant to added DNAase

It is likely that rDNA cannot cross the cell membrane because of its bulky adducts (rhodamine plus a positively charged linker) and therefore accumulates in the periplasm. Although we have found that rDNA with a labeling density similar to that used in this study is reduced in its ability to transform, two considerations make it impossible to interpret this reduced transforming capacity. First, low density random labeling will leave variable stretches of label-free DNA that may be capable of transforming. On the other hand, transformation could be blocked by the adducts at the levels of recombination, gene expression or replication.

To investigate the location of DNAase-resistant rDNA further, we generated 3D-deconvolved, volume-reconstructed images of transformed CFP- and YFP-labeled competent cells with and without DNAase treatment (Figure 1C and D). The images show representative cells, viewed from 4 directions, related by 90° rotations around the long axis of the cell. The three images in Figure 1C show a typical cell not treated with DNAase and imaged after 2 minutes of incubation with rDNA, a time when very little tDNA is protected from DNAase. The projected views show a blob of rDNA, exhibiting limited contact with the surface of the YFP volume. λ DNA molecules in solution are coils with an average radius of gyration equal to about 500 nm (41). Figure 1C is consistent with binding of such an rDNA coil to the cell surface, involving a limited portion of the rDNA surface. In cells imaged after 30 minutes of incubation, whether DNAase-treated or not, the rDNA is more closely juxtaposed to the surface (Figure 1D). In Figure 1D some of the rhodamine signal is stretched across the surface, and comparison of the rotated views shows rDNA that appears to wrap around the cell volume, consistent with its location within the confined space of the periplasm and its extension across the outer face of the cell membrane. Figure 1E shows a single optical slice through three cells, imaged after DNAase treatment following 30-minutes incubation with rDNA, confirming that the rDNA signal is localized near the edges of the cells, consistent with a periplasmic location and with the failure of rDNA to cross the membrane. Figure 1C-E thus suggests a process in which rDNA initially associates with the wall surface by localized contact and then enters the periplasmic compartment where it becomes DNAase-resistant, while some of it then extends across the cell membrane surface.

### ComFA stabilizes ComGA

Before studying the roles of individual proteins in rDNA uptake, it was important to determine the impact of each deletion mutation on the stabilities of other proteins needed for transformation. Figure S1 shows Western blots in which we have examined Δ*comFA, ΔcomGA, ΔnucA, ΔcomC, ΔcomEA* and *comEC518* loss-of-function mutant extracts using antisera raised against ComGA, ComFA, NucA, ComEA and ComEC. All the mutations are deletions except that *comEC518* is a Tn917 insertion, which exhibits no transformability and is located near the start of the *comEC* coding sequence after residue V50. All of the proteins are present at close to normal levels in the mutants, with the important exception of ComGA, which is destabilized in the absence of ComFA (Figure S1B), as reported previously (42) and suggesting that ComGA and ComFA are binding partners. However, ComGA is present at the normal level in an inactivating K152E mutant of *comFA* (33, 35). K152 lies in the Walker A motif of ComFA, demonstrating that the ATPase activity of ComFA is not needed for ComGA stability. In fact, ComGA interacts with both ComFA and ComFC (43). Thus, any uptake-associated Δ*comFA* phenotype may be ascribed to an indirect effect on ComGA

### Testing mutants for rDNA binding and uptake

We next examined mixtures of YFP- and CFP-labeled wild-type and mutant strains. To analyze the raw images, we utilized several tools. First, we counted the percent of competent cells for mutant and wild-type strains that were associated with detectable rDNA signals. Second, when required, we recorded deconvolved images and performed volume reconstructions. These separate types of measurement were helpful because not all the competent cells were associated with rDNA, even in the wild-type and because in theory a given mutant might not have 100% penetrance, so that a few cells might exhibit relatively normal binding or uptake while others are totally deficient. It is important to recognize that the images recorded without DNAase treatment reflect rDNA binding plus uptake, whereas after treatment they reflect uptake to the periplasm.

### ComGA and ComC

The Δ*comGA* and Δ*comC* mutants exhibited no detectable association with rDNA even without DNAase treatment (Figure S2). Because ComGA is needed to assemble tpili and ComC is a peptidase that processes the major and minor pilins for assembly (23), these results support the hypothesis that tDNA binding requires tpili, most likely because the DNA binds directly to these organelles, as it does in *S. pneumoniae* (20) and in *V. cholerae* (6).

### ComFA

In the Δ*comFA* strain, there is a significant decrease in the percent of cells that were associated with rDNA after 30-minutes, either with or without DNAase treatment (Table 1). Despite this, many Δ*comFA* cells did bind and take up rDNA. In the absence of DNAase treatment the signal intensities of the wild-type and mutant cells were similar (Figure S3, panel A) but with treatment the signals in the mutant were markedly weaker (Figure S3, panel B), in agreement with results obtained using radiolabeled tDNA (32, 34). If the decrease in signal intensity in the Δ*comFA* strain is due to a deficiency in ComGA (Figure S1, panel B), we would expect the *comFA* K152E mutant to behave like the wild-type. Indeed, no significant effect of the K152E mutant was observed on the percent of cells showing rDNA fluorescence either with or without DNAase treatment (Table 1) and the images in Figure 2A and 2B show that there are no obvious effects of the mutation on either the total or DNAase resistant signal intensities. We conclude that the effect of Δ*comFA* on the acquisition of DNAase resistance and on rDNA binding to the cells is largely due to ComGA deficiency. Despite the non-requirement of ComFA for uptake, the rDNA signal in the K152E mutant cells is more localized after DNAase treatment than in the wild-type cells, suggesting an unexpected ATP-dependent requirement for ComFA in the spreading of the rDNA in the periplasm (Figure 2C).

**Figure 2.**
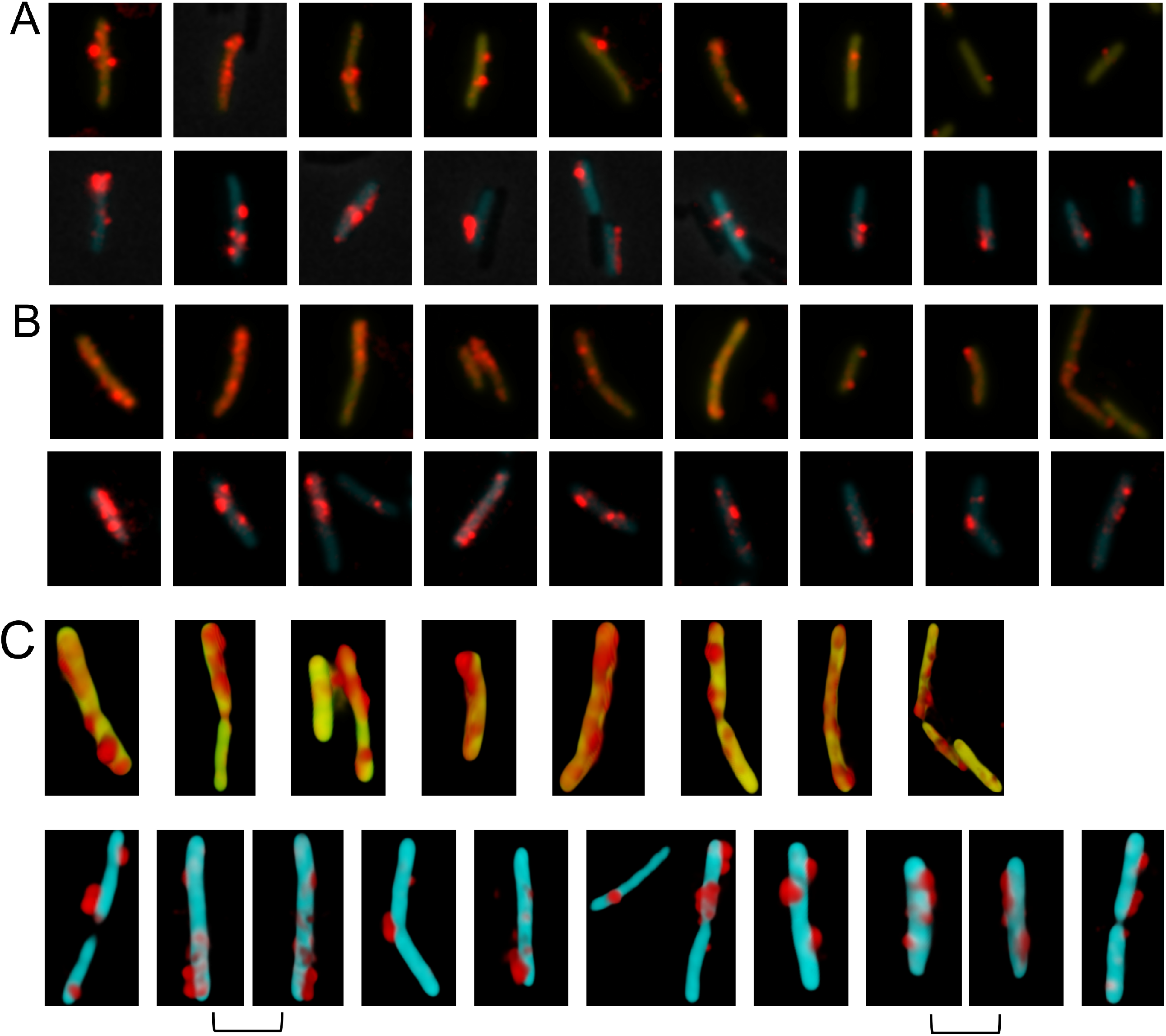
Binding and uptake in the *comFA* K152A mutant. Wild-type (YFP) and mutant (CFP) cells were combined before 30 minutes incubation with rDNA. Panels A and B show results for *comFA* K152E without and with DNAase treatment respectively. In each panel the top and bottom rows show 9 each of wild-type and mutant cells respectively. All the cells were taken from a single representative microscope field and show all or nearly all of the rDNA-associated cells in that field. In the cropped images of a given panel, rDNA signals were enhanced identically but rDNA signal intensities cannot be compared between panels A and B. Panel C shows volume reconstructed images of several representative wild type (top row) and mutant (bottom row) cells. In each case a single view is shown except for the adjacent images joined by brackets, which show two views of the same cell related by 180° rotations.

**Table 1.**
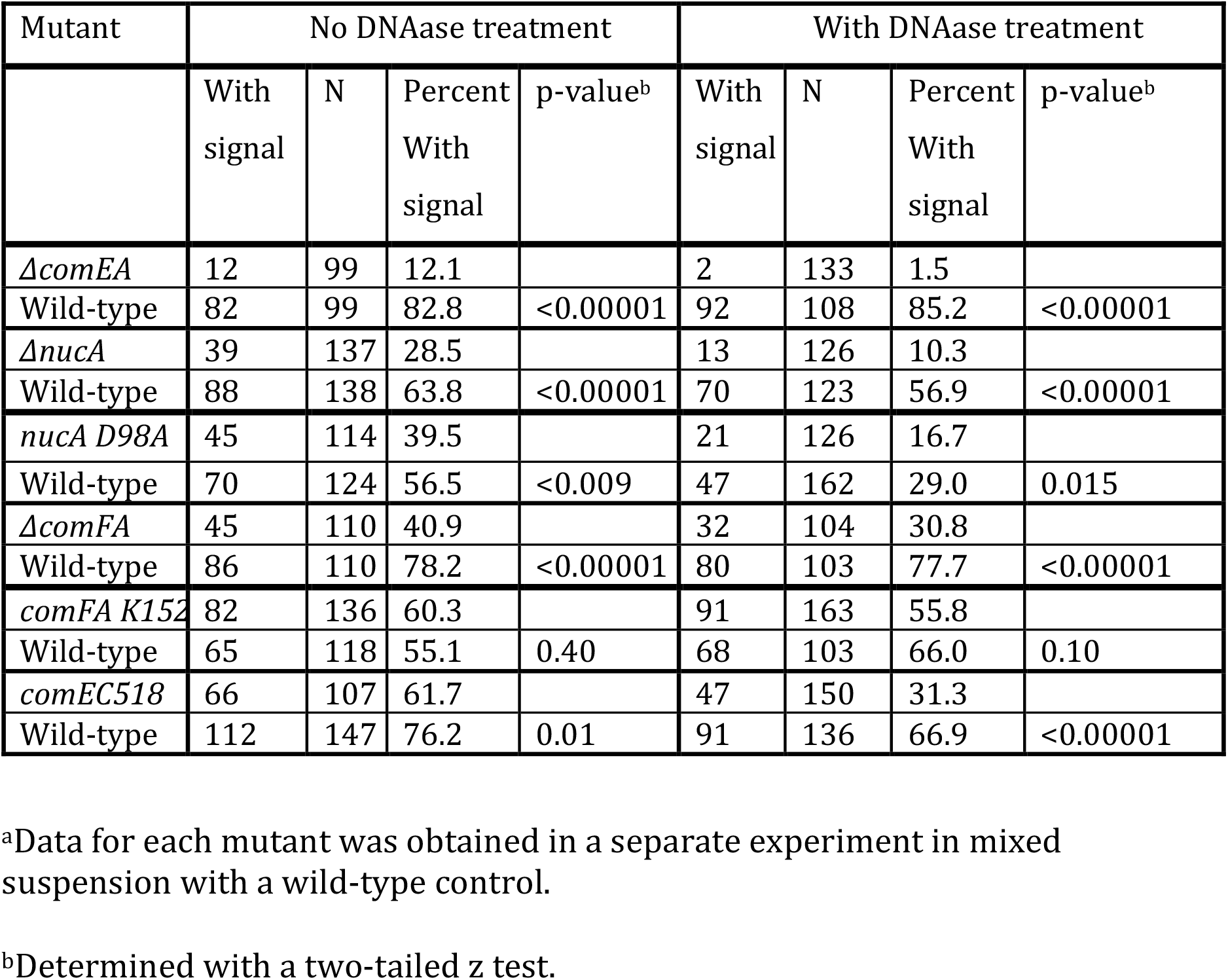
Binding and uptake of rDNA by transformation mutants^a^

| Mutant | No DNAase treatment |  |  |  | With DNAase treatment |  |  |  |
| --- | --- | --- | --- | --- | --- | --- | --- | --- |
|  | With signal | N | Percent With signal | p-value<sup>b</sup> | With signal | N | Percent With signal | p-value<sup>b</sup> |
| *ΔcomEA* | 12 | 99 | 12.1 |  | 2 | 133 | 1.5 |  |
| Wild-type | 82 | 99 | 82.8 | <0.00001 | 92 | 108 | 85.2 | <0.00001 |
| *ΔnucA* | 39 | 137 | 28.5 |  | 13 | 126 | 10.3 |  |
| Wild-type | 88 | 138 | 63.8 | <0.00001 | 70 | 123 | 56.9 | <0.00001 |
| *nucA D98A* | 45 | 114 | 39.5 |  | 21 | 126 | 16.7 |  |
| Wild-type | 70 | 124 | 56.5 | <0.009 | 47 | 162 | 29.0 | 0.015 |
| *ΔcomFA* | 45 | 110 | 40.9 |  | 32 | 104 | 30.8 |  |
| Wild-type | 86 | 110 | 78.2 | <0.00001 | 80 | 103 | 77.7 | <0.00001 |
| *comFA K152* | 82 | 136 | 60.3 |  | 91 | 163 | 55.8 |  |
| Wild-type | 65 | 118 | 55.1 | 0.40 | 68 | 103 | 66.0 | 0.10 |
| *comEC518* | 66 | 107 | 61.7 |  | 47 | 150 | 31.3 |  |
| Wild-type | 112 | 147 | 76.2 | 0.01 | 91 | 136 | 66.9 | <0.00001 |

<sup>a</sup>Data for each mutant was obtained in a separate experiment in mixed suspension with a wild-type control.
<sup>b</sup>Determined with a two-tailed z test.

### ComEA

In a Δ*comEA* strain the percent of cells with associated rDNA signals was reduced about 7-fold in the absence of DNAase treatment (Figure 3A and Table 1) and the frequency of cells with DNAase resistant rDNA was reduced 57-fold (Figure 3B and Table 1), consistent with results obtained with *B. subtilis* and *S. pneumoniae* using radiolabeled DNA (19, 37, 38). In the Δ*comEA* cells that did exhibit binding, the rDNA remained on the cell surface (Figure 3C), like the wild-type image at 2-minutes (Figure 1A). Thus, ComEA assists in binding and is required for uptake. Most likely, the initial binding of tDNA to the tpilus is labile and must be stabilized by contact with periplasmic ComEA, a known DNA-binding protein. Indeed, reversible attachment of DNA to the surface of *B. subtilis* competent cells has been reported (44). The nearly total DNAase susceptibility of rDNA in the mutant demonstrates that ComEA is also needed for uptake to the periplasm.

**Figure 3.**
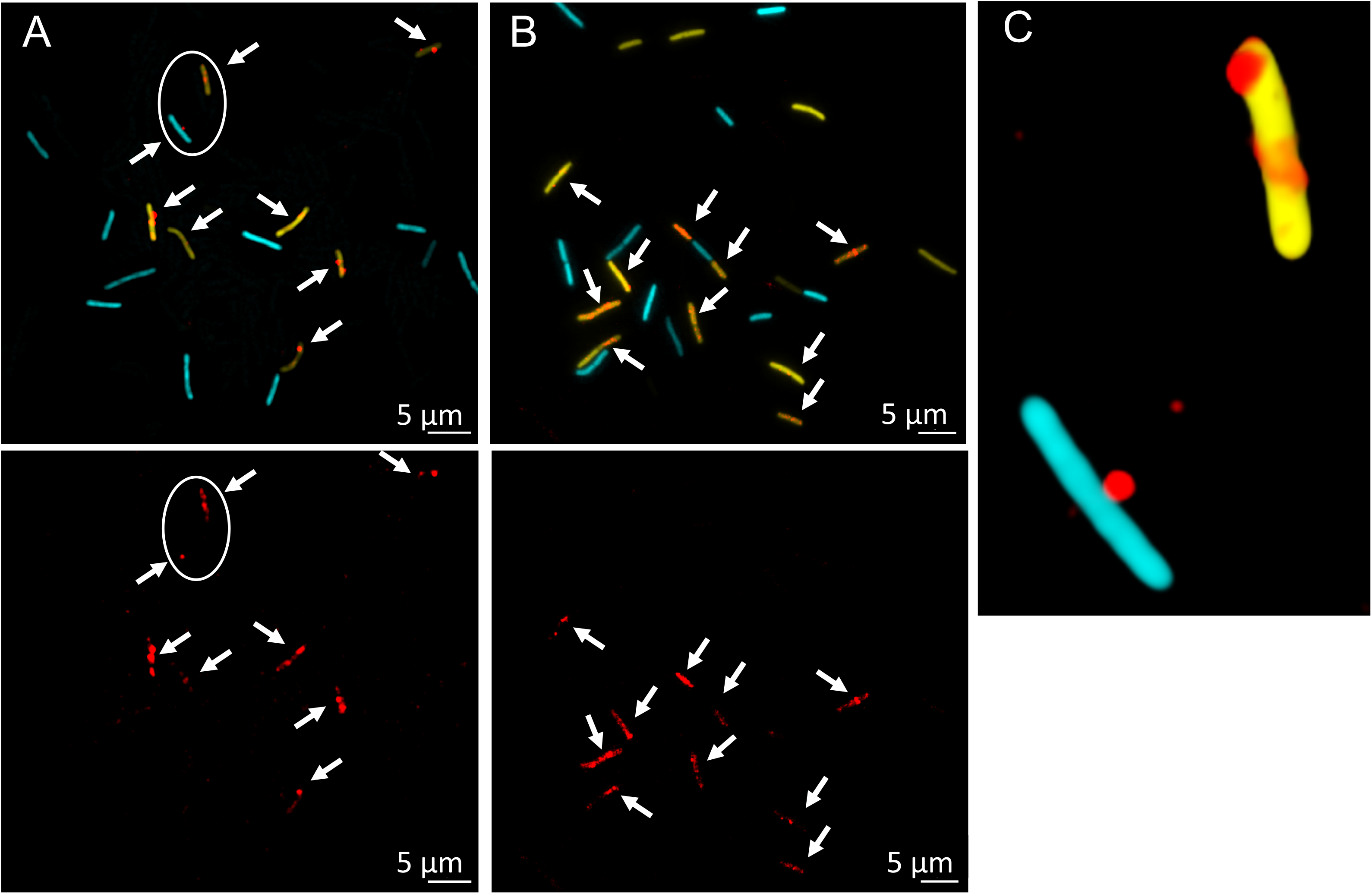
Binding and uptake in the Δ*comEA* mutant. Wild-type (YFP) and Δ*comEA* cells were combined before incubation for 30-minutes with rDNA. Panel A shows the cells without DNAase treatment and B shows cells with treatment. In the top images the rhodamine, CFP and YFP channels were merged and the bottom images in A and B show only the rhodamine channel. The arrows indicate the position of all the rhodamine signals. Panel C shows one aspect from a 3D deconvolution of the CFP- and YFP-expressing cells circled in Panel A.

### NucA

A Δ*nucA* strain exhibits decreased rates of transformation, tDNA binding and the acquisition of DNAase resistance (37). To explore this phenotype with rDNA, we used samples incubated with rDNA for 15-rather than 30-minutes to maximize the differences between the mutant and wild-type strains, because the final yield of transformants in the mutant is only reduced about 2-fold (37). The percent of Δ*nucA* cells associated with rDNA (Table 1) was reduced about 2-fold and the rDNA signal intensities were clearly reduced after DNAase treatment (Figure S4). To determine if the uptake deficiency of the Δ*nucA* strain was due to the nuclease activity of NucA, we inserted a D98A active site mutation at the native *nucA* locus. The design of this mutation was based on analysis of NucB (45), which exhibits 58% identity with NucA. D98A corresponds in position to residue D87A in NucB. This mutant exhibits a transformation deficiency similar to that of the Δ*nucA* strain. The data in Table 1 show that the D98A strain has a similar phenotype to the knockout of *nucA*, although the effect on the frequency of cells associated with rDNA is not as great, perhaps due to residual nuclease activity. Figure 4 shows that the intensities of the rDNA signals in this mutant were decreased following DNAase treatment, confirming that the nuclease activity of NucA is needed for uptake to the periplasm and excluding a polar effect of *nucA* inactivation on the downstream *nin* gene (46). Volume reconstructions after DNAase treatment for the Δ*nucA* (Figure S5 A) and D98A (Figure S5 B) mutants show that the rDNA is more localized than in the wild-type. similar to the phenotypes of the *comFA* K152E and *comEC518* mutants (see below). Based on the earlier assumption that DNAase resistance corresponds to entry to the cytoplasm, it was proposed that NucA produced ends for transport through the membrane channel (37). Although this may be true, the present results show that the nuclease activity also plays a role in uptake to the periplasm.

**Figure 4.**
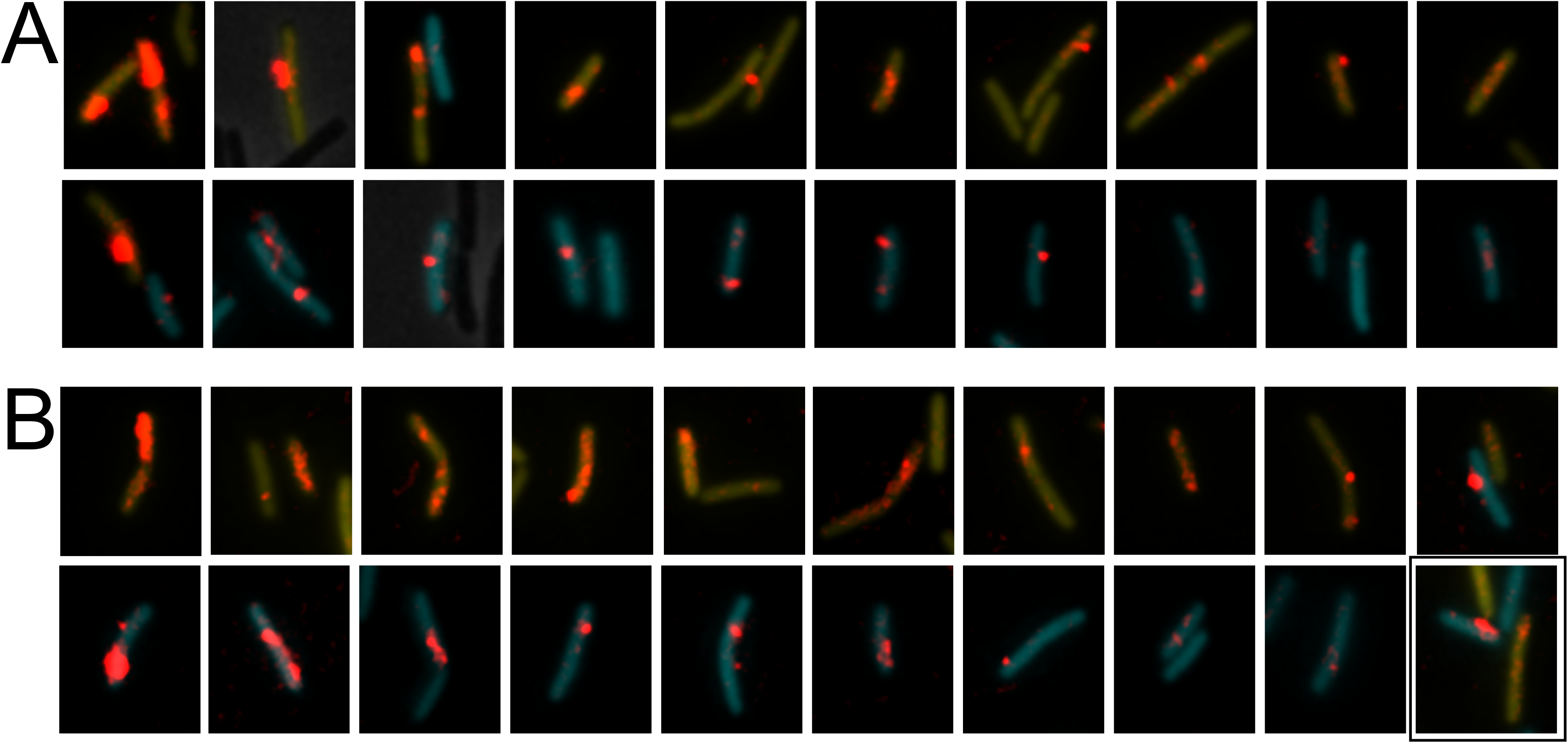
Binding and uptake in the *nucA* D98A mutant. Wild-type (YFP) and D98A (CFP) mutant cells were made competent, combined, and after 15 minutes incubation with rDNA were imaged without (panel A) and with DNAase (panel B). In each of these panels the top and bottom rows show nearly all the wild-type and mutant cells from a single field, respectively. The rhodamine signal was enhanced before cropping the cells and the intensities of the rhodamine signal can be directly compared. The YFP and CFP images were separately adjusted in each image so as not to obscure the rDNA signal. The images in these two panels were ordered by apparent rDNA signal strengths decreasing from left to right, to facilitate comparisons of the mutant and wild-type cells. The boxed image in panel B shows both a YFP- and a CFP-expressing cell.

### ComEC

As noted above, the use of radiolabeled rDNA showed that the elimination of ComEC did not affect the total association of tDNA with competent cells but markedly reduced the label after DNAase treatment (4, 37, 38). The use of rDNA should therefore show a previously unexpected role for ComEC in uptake. To test this, we used the *comEC518* mutation, which has a transposon insertion near the N-terminus of the *comEC* coding sequence (47). After 30 minutes, the fraction of competent cells showing DNAase-resistant rDNA was moderately decreased in the *comEC518* mutant (31%) compared with 67% for the wild-type (Table 1), probably a significant difference (p=0.01). In contrast, there is little if any effect on the frequency of rDNA attachment before DNAase treatment. This trend is confirmed by the images contained in Figure 5. Panel A shows that in the absence of DNAase treatment, the intensities of the rDNA signals are comparable in wild-type and mutant cells. However, Figure 5B shows that the rDNA signal in those DNAase-treated *comEC518* cells that exhibit uptake is markedly lower than in the wild-type, in agreement with measurements made using radiolabeled tDNA (4, 19, 34, 37). Interestingly, the volume reconstructed images in Figure 5C reveal a marked difference between the DNAase-resistant rDNA signals in wild-type and *comEC* cells. In the wild-type cells, as noted above (Figure 1B), some of the rDNA is extended across the volume surface. But in the *comEC518* cells, the DNAase resistant rDNA is more localized. Thus, although binding to the cell surface is essentially normal in the *comEC* mutant, uptake is strongly affected and the reduced amount of rDNA that manages to enter the periplasm is confined locally.

**Figure 5.**
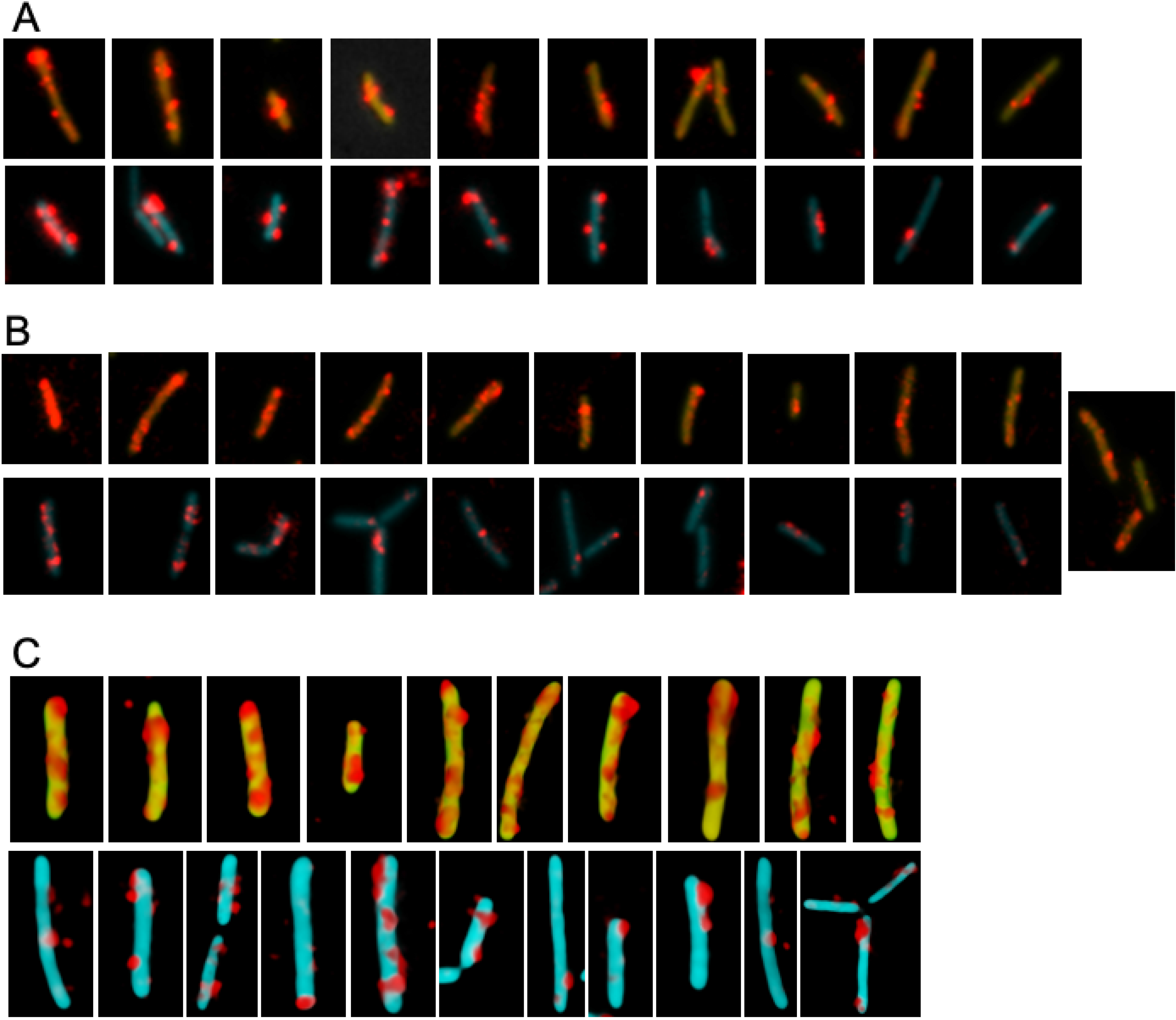
Binding and uptake in the *comEC518* mutant. Wild-type (YFP) and Δ*comEC518* (CFP) cells were combined before 30-minutes incubation with rDNA. The images were processed as described in the legend to Figure 4. Panel A shows the cells without DNAase treatment and B shows cells with treatment. The single image to the right of panel B shows two YFP cells and one CFP cell. The images in these two panels were ordered by apparent rDNA signal strengths decreasing from left to right, to facilitate comparisons of the mutant and wild-type cells. Panel C shows selected volume reconstructed images of wild-type and *comEC518* cells selected from the same field, representing almost all the cells in the field. Only one view is shown of each cell. Image intensities cannot be compared either within panel C or with the images in panels A and B.

### rDNA closely associates with YFP-ComEA during uptake

In the Gram-negative systems, periplasmic ComEA is initially distributed uniformly within the periplasm and then dramatically forms foci, accumulating at the sites of DNA uptake (8, 9). To examine the interactions of rDNA with ComEA *in vivo*, we constructed a YFP-ComEA mutant in which the N-terminal YFP moiety was in the cytosol and the DNA binding portion of ComEA was appropriately located in the periplasm. The *yfp-comEA* construct was placed at the *comE* locus, expressed normally under competence control. The resulting strain, with *yfp-comEA* as the only source of *comEA* in the cell, was normally transformable, although the rate of appearance of DNAase resistant transformants was deceased about 2-fold. As reported previously using immunofluorescence with native protein (48) and by Kaufenstein et al (49) with a YFP-ComEA construct similar to ours, ComEA forms large foci at several locations around the membrane. Uniquely among the transformation proteins there is no obvious preference of these foci for polar or sub-polar locations. The foci are not caused by the uptake of DNA from occasional lysed cells, because the addition of DNAase (100 μg/ml) during 90 minutes of growth prior to sampling had no effect on this distribution.

After the addition of rDNA, the distribution of YFP-ComEA foci was not obviously different in the cells with and without bound rDNA. Figure 6A shows deconvolved and volume reconstructed images of four representative cells from a single field that exhibit significant rDNA attachment after I minute incubation with rDNA. In these images, the rhodamine signal is almost always localized near the poles as reported previously using a different DNA-labeling protocol (48). The rDNA in Figure 6A, is consistently near a blob of YFP-ComEA in all the cells shown except in panel Aa, which may show a cell with initial attachment to a tpilus that is not located near an accumulation of ComEA. In the remaining cells of panel A, the rDNA and YFP-ComEA appear to be connected by faint regions of YFP signal intensity. After 10 minutes of incubation the images are strikingly different (Figure 6B). At this time, the rDNA is often intimately associated with foci of YFP-ComEA and is sometimes located away from the cell poles (Figure 6B, cells c and d). At this time, in this same experiment, much of the rDNA intensity is resistant to DNAase. These images are consistent with a scenario in which tDNA initially attaches to tpili at the poles, and is then pulled into the periplasm, where it binds to locally mobile molecules of ComEA. The Graumann laboratory has reported that the diffusive behavior of YFP-ComEA in the membrane is similar to that of tDNA in the periplasm, suggesting that these molecules are associated (49), consistent with this conclusion. After the capture event, which results in stabilized binding but no major redistribution of ComEA, the tDNA is plausibly pulled into the periplasm by a Brownian ratchet mechanism involving ComEA and ComEC, where it is stretched across clusters of ComEA molecules.

**Figure 6.**
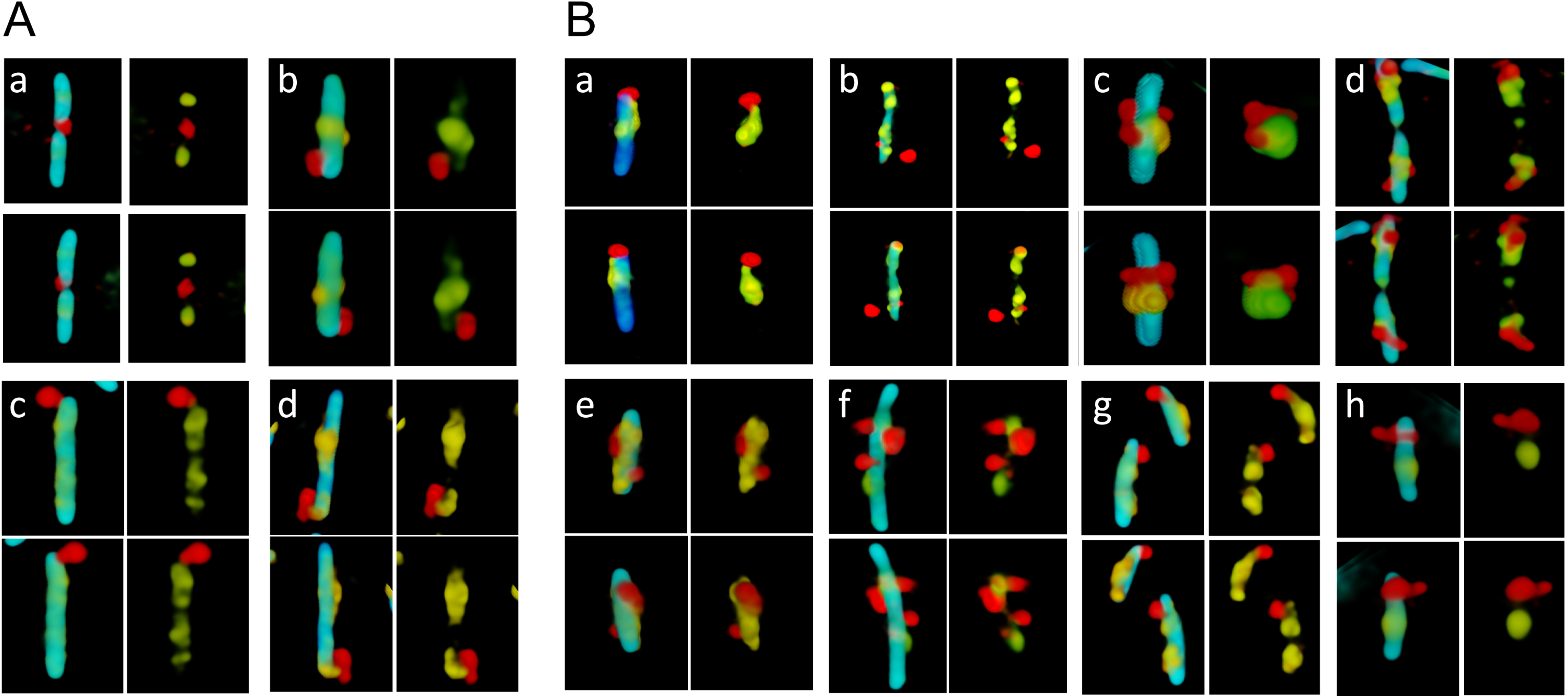
Association of rDNA with YFP-ComEA without DNAase treatment. In each group of four images (a-d in panel A and a-h in panel B), the top and bottom pairs show views of a single cell, related by 180° rotations. The left-hand images in each group show the superimposed YFP, rhodamine and CFP channels, where CFP delineates the cytoplasmic volume. The right-hand images show only the YFP and rhodamine channels, for clarity. Panel A shows images collected after 1 minute incubation of the cells with rDNA and Panel B after 10 minutes.

## DISCUSSION

The new appreciation of the meaning of DNAase resistance allows us to interpret data acquired in this study and to reinterpret data from the literature. It had been generally assumed that the acquisition of resistance was due to transport of tDNA across the cell membrane, but our present results show that resistance is actually imparted by the cell wall barrier. DNAase, has a calculated molecular weight of 29 kDa and it has been estimated from *in* vitro studies that globular proteins of more than about 25 kDa cannot diffuse through the cell wall of *B. subtilis* (50). The wall preparations used to produce this estimate were treated to remove associated proteins and the intact wall is likely to be even denser and thus more restrictive to diffusion.

The conceptual partition of transformation proteins into three exclusive categories based on their requirements for binding, uptake and transport, does not fit with the new data. Thus, ComEA is needed for both tight binding and uptake. ComGA appears to be needed for both initial binding and for uptake because the Δ*comFA* mutation, which lowers the amount of ComGA, decreases the frequency of rDNA binding (Table 1) and is deficient in the amount of uptake in individual cells (Figure S3), while the *comFA* K152E mutation does not reduce uptake. NucA, ComEC and ComFA help to distribute tDNA within the periplasm and the latter two proteins are certainly also needed for transport to the cytoplasm, while NucA helps with binding as well as uptake. This complexity is consistent with the participation of the proteins in a nanomachine, in which they interact to guide tDNA through the wall, periplasm and membrane. Further support for this notion is provided by the observations that ComFA stabilizes ComGA (Fig, S1) and that ComFC binds to ComFA (31). Co-immunoprecipitation experiments in *B. subtilis* have shown that ComGA associates with ComFA, ComFC, DprA, SsbB, RecA and ComEC (43), all proteins that participate in transformation, providing support for the notion of a nanomachine.

The results for Δ*comGA* and Δ*comC* conform to previous data for *B. subtilis* and *S. pneumoniae* obtained using radiolabeled DNA (24, 44, 51) showing that in the absence of these proteins there is no detectable binding of tDNA to the mutant cells. This reinforces the belief that the tpilus is the initial site of tDNA binding and is consistent with the localization of both ComGA and the initial sites of DNA binding near the poles (48). tpilus retraction has now been observed in both *V. cholerae* and *S. pneumoniae* (6, 22), and is very likely to also occur in *B. subtilis*, although both of these Gram-positive organisms lack retraction ATPases. Based on a number of recent studies it is tempting to conclude that the *B. subtilis* pilus retracts by spontaneous disassembly into the membrane without an external energy source (6, 52–54). However, a dual function ATPase has been described that drives both assembly and retraction of a t4 pilus (55), and this remains a possibility for ComGA. In fact, the major effect of Δ*comFA* on uptake compared to binding (Table 1, Figure S3), could be explained by the decreased amount of ComGA (Figure S1),

In the absence of ComEA, tDNA binding is reduced, and uptake is eliminated. There is experimental evidence for an initially transient form of binding (44), suggesting that attachment of tDNA to the tpilus is reversible. ComEA is too small to extend from the membrane through the wall to aid in the initial binding, and attachment of tDNA to the cell is likely to be stabilized by contact between the tDNA and ComEA in the periplasm. During the uptake of tDNA, no dramatic relocation of the membrane-anchored YFP-ComEA comparable to the reorganization reported for *Vibrio* and *Neisseria* (8, 9, 12) was observed. We propose that inwardly diffusing DNA segments are first captured by fixed or locally mobile ComEA molecules. Further uptake and the spreading of rDNA within the periplasm then results from the inward diffusion of additional segments of DNA, which are trapped by more distant ComEA molecules, in an on-going Brownian ratchet process. The dependence of uptake on NucA and ComEC and the restricted distribution of DNAase resistant rDNA in the *comEC518, nucA* and *comFA* K152E mutants suggests that a more complex mechanism takes place than in the Gram-negatives, where uptake depends only on the tpilus and ComEA. This complexity may be characteristic of bacteria with membrane-anchored, relatively immobile ComEA.

A role for ComEC in uptake to the periplasm was unexpected. The *B. subtilis* ComEC contains two conserved periplasmic domains that are obvious candidates for involvement in uptake (29, 56, 57): the N-terminal N-loop or OB-domain (PFAM13567) and the C-terminal metallo-β-lactamase domain (PFAM00753) (57). Between the two is the competence domain (PFAM03772) that contains several transmembrane helices and must contribute to forming the transport channel. The restricted localization of rDNA in *comEC* and *comFA* <u>mutants</u> suggest that tDNA is transferred to ComEC in a process that depends on ComFA. The absence of ComEC might then arrest uptake by confining the rDNA to locally available ComEA. Transfer to ComEC would also position tDNA for entry to the channel and for degradation of the non-transforming strand. A recent finding in *Helicobacter pylori* (58) is consistent with these ideas. This organism encodes neither tpili nor ComEA but uses a type 4 secretion system to bring tDNA into the periplasm and ComH, a periplasmic DNA binding protein that interacts with tDNA in the periplasm. Suggestively, ComH, which does not resemble ComEA but plays an analogous role, interacts directly with the OB-domain of ComEC (58). It has been proposed that the β-lactamase domain contains the nuclease activity that degrades the non-transforming strand of tDNA (57). If degradation does take place in the periplasm, ComEC may provide an entropic boost to uptake by converting tDNA to single strands and free nucleotides.

The role of NucA in uptake is also a novel finding. NucA is anchored in the membrane with its catalytic site protruding into the periplasm. Its nuclease activity assists in uptake (Table 1, Figure 4) implying that either a nick on one strand of tDNA or a double strand cut facilitate uptake. Indeed, a double strand cut is likely, as suggested by studies of the NucA paralog, NucB (57). Transformation with circular DNA has shown NucA-dependent conversion to the full length linear form shortly after binding (37). This result reinforces the idea that tDNA crosses the wall as a folded dsDNA molecule. Folding DNA, which has a persistence length of about 50 nm (~147 bp), requires local deformation of the DNA structure, presumably caused by binding to the tpilus. After release from the tpilus, strain in the folded tDNA would impel the tDNA to straighten locally and pull back through the wall. A double strand break induced by NucA would relieve the strain and accelerate the ratchet. An additional explanation for the role of NucA, which is not mutually exclusive with this one, is suggested by the finding that the ComEA protein of *V. cholerae* prefers binding to a DNA end (8). If this is also true of the *B. subtilis* protein, NucA might not only relieve the strain in folded tDNA but would also facilitate the initial binding of ComEA to the incoming DNA. These ideas are consistent with our finding that both binding and uptake are reduced in the *nucA* mutants (Table 1, Figure 4). The termini produced by NucA would also be available to enter the ComEC channel (37).

The results presented in this study allow us to propose an updated model for transformation in *B. subtilis*. First, tDNA binds reversibly to the tpilus, which retracts, bringing a folded loop of DNA into the periplasm. The *S. pneumoniae* tpilus is 6 nm thick and retraction would likely leave a hole in the wall of the same size. A folded DNA molecule would have thickness of 4 nm, permitting the tpilus to bring the DNA into the periplasm with its hydration shell intact, and allowing subsequent inward diffusion of the DNA. NucA then cuts the tDNA as it is released from the tpilus, relieving strain introduced by tDNA folding and possibly facilitating binding to ComEA at a terminus. This binding stabilizes the association of tDNA with the cell and as successive segments of DNA diffuse into the periplasm through the cell wall, they are captured by additional ComEA molecules and prevented from diffusing outward. As the ComEA binding sites near the locus of entry become occupied, tDNA spreads in a process dependent on ComFA and ComEC and is captured by ComEC for transport to the cytoplasm, accomplished by ComEC and the ComFA/ComFC complex, which comprise the transformation permease.

The use of labeled DNA that cannot cross the membrane may obscure what is actually a seamless process of uptake and transport. In other words, transformation may not be a two-step process in which tDNA accumulates in the periplasm. Another open question concerns the extent to which our revised understanding of DNAase resistance applies to other Gram-positive transformable bacteria, such as *S. pneumoniae*. Despite the shared requirements for ComEA, ComEC and t4 pilus proteins, there are likely to be important mechanistic differences between the uptake mechanisms of these two model Gram-positive bacteria. In *S. pneumoniae*, degradation of the non-transforming strand is accomplished by EndA (59, 60) and degradation takes place in the absence of ComEC (38), implying that the latter is not needed for uptake to the periplasm. In contrast, *B. subtilis* does not encode EndA and when *comEC* is inactivated, degradation ceases (37), consistent with the role of ComEC in uptake and perhaps with its proposed action as a nuclease. Finally, in *B. subtilis* the tpilus is probably short while in *S. pneumoniae* it extends from the cell surface (20, 23). Many outstanding questions remain, most importantly, how these various processes are accomplished on the molecular level.

## Materials and Methods

### Strains and growth conditions

All mutant strains (Table S1) of *B. subtilis* were constructed in IS75 (his leu met) and are derivatives of the domesticated strain 168. All the strains expressed YFP or CFP controlled by the *comG* promoter, inserted ectopically in *amyE. B. subtilis* was grown to competence for all experiments by the two-step method (61) except that unfrozen cultures were used and cultures were started from plates grown overnight instead of from spore suspensions.

### Transformation for microscopy

Competent cultures of P_G_-CFP and P_G_-YFP expressing mutant and wild-type strains were mixed 1:1 and the mixture was incubated with 0.2 μg/ml of rhodamine-labeled DNA for the indicated times, in final volumes of 300-500 μl. Samples were taken and either immediately fixed by the addition of 3.2% paraformaldehyde and incubation at room temperature for 30 minutes or were first incubated for 3 minutes with 100 μg/ml DNAase I at 37° C before fixation. In the experiment shown in Figure 1, where DNAase-treated and untreated samples were combined for visualization, the cells were centrifuged briefly and resuspended in the same medium used to achieve competence, but containing no Mg and EDTA (10 mM) to stop the nuclease activity before mixing and fixation. Samples were then washed and imaged.

### Preparation of rDNA

Bacteriophage λ DNA (New England Biolabs) was diluted to 10 μg/ml and incubated for 60 minutes with a MIRUS Label-IT rhodamine TM reagent (Mirus Bio) according to the manufacturer’s recommendations. The DNA was then processed through a spin column to remove excess reagent.

### Point mutation of *nucA*

A fragment of DNA carrying the *nucA* sequence with a D98A mutation and *Bam*H1 ends, was synthesized by Biomatik (Canada) and cloned into the BamH1 site of pMiniMAD2, a gift from Dan Kearns (Indiana University). After transformation with selection for erythromycin (5 μg/ml), the markerless chromosomal mutant was isolated as described by Patrick and Kearns (62).

### YFP-ComEA construction

Three fragments were produced by PCR and assembled using NEBuilder HiFi into pUC18CM, a derivative of pUC18 with a chloramphenicol resistance cassette for selection in *B. subtilis.*. Fragment #1 carried 1500 base pairs upstream from the *comEA* coding sequence including the promoter, the ribosomal binding site and start codon. To produce this fragment primers 1 and 2 were used (Table S2). Fragment #2 carried the*yfp* open reading frame with the stop codon omitted, produced using primers 3 and 4. Fragment #3 consisted of the *comEA* open reading frame without its start codon, produced using primers 5 and 6. After assembly, the clone was verified by sequencing and transformed into the native locus via a single crossover event. The duplicated native *comEA* open reading frame was then deleted by transformation using DNA from strain BD8739 *(comEA::ery trpC2)* and the final construction was again verified by sequencing.

### Microscopy

For microscopy 1 μl samples of transformed cells were placed on thin agarose pads before visualization. All images were acquired with a 100x Plan Appo immersion objective, NA=1.40 on a Nikon T*i* microscope, equipped with LED excitation sources and an Orca Flash 4.0 camera (Hamamatsu). Nikon Elements was used for data acquisition and image analysis. Z-stack images were recorded with 200 nm steps and processed for 3D deconvolution with the Landweber algorithm in Nikon Elements. Volume reconstructions were assembled with alpha blending. Images were exported to Adobe Photoshop and Microsoft PowerPoint for final processing.

### Western blotting

Western blotting was carried out using standard methods with semi-dry blotting and the nitrocellulose blots were developed using ECL (GE Healthcare). The images were recorded with a BioRad ChemiDoc MP imager. In most gels, for a loading control, the top of the membrane was cut off after blotting and developed separately with anti-EFG antiserum, a kind gift from Jonathan Dworkin (Columbia University Medical school).

### Statistical analysis

Proportions of mutant and wild-type cells that were associated with rDNA were compared using a two-tailed z-test.

## Data availability statement

The data that support the findings of this study are available from the corresponding author, [author initials], upon reasonable request.

## SUPPLEMENTAL MATERIAL

Table S1 Strains

Table S2 Primers

Figure S1. Western blots for transformation proteins in mutant strains.

Figure S2. DNA binding in Δ*comGA* and Δ*comC* strains.

Figure S3. Binding and uptake in the Δ*comFA* mutant.

Figure S4. Binding and uptake in the Δ*nucA* mutant.

Figure S5. Volume reconstructions of the Δ*nucA* and *nucA* D98A mutant strains.

## ACKNOWLEDGEMENTS

This work was supported by NIH grant R01GM057720. We thank Mathew Neiditch and members of the Dubnau lab for helpful comments and discussion. We thank Daniel Ziegler at the Bacillus Genetic Stock Center for providing strains and J. Dworkin for the anti-EFG antiserum. J. H. and M. D. S. performed all of the experimental work. D. D. supervised the work and all the authors contributed to planning experiments, analyzing data and to writing the paper.

---

**Table S1.**
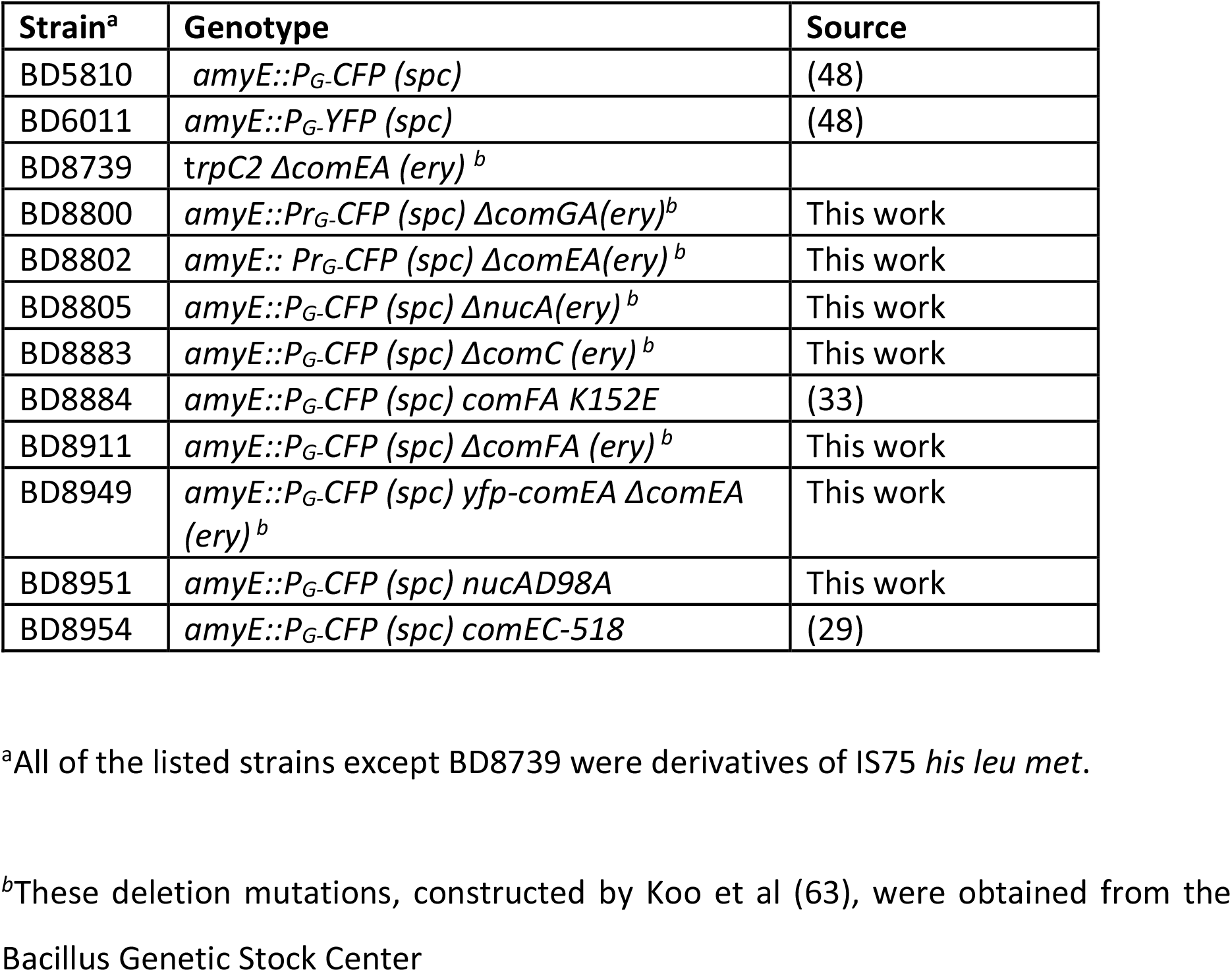
Strains

**Table S2.**
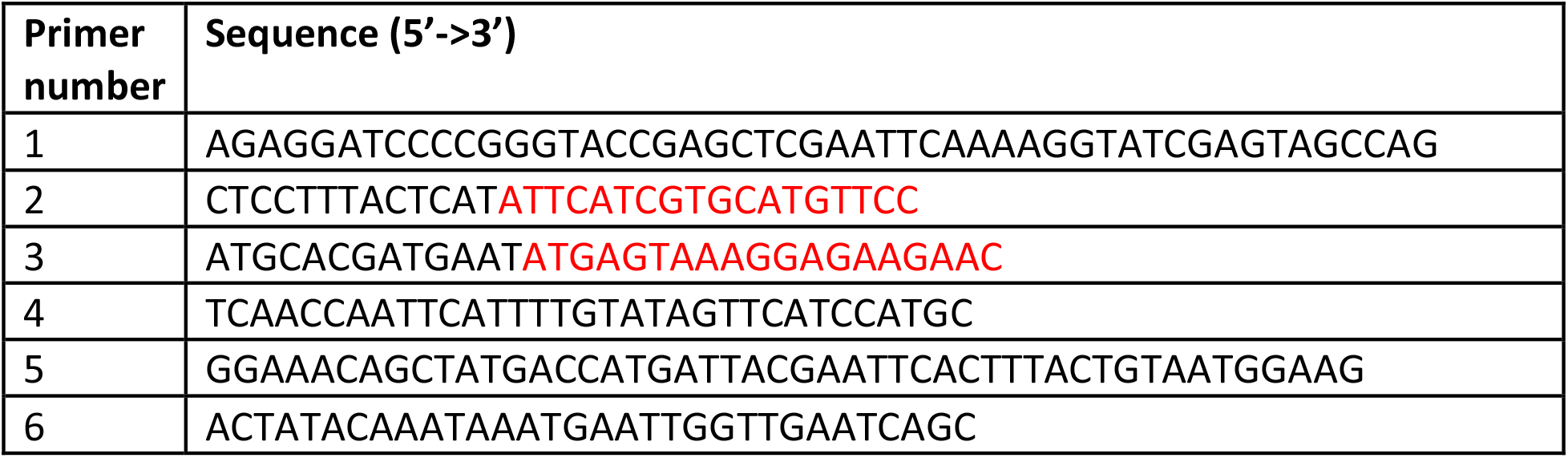
Primers

**Figure S1.**
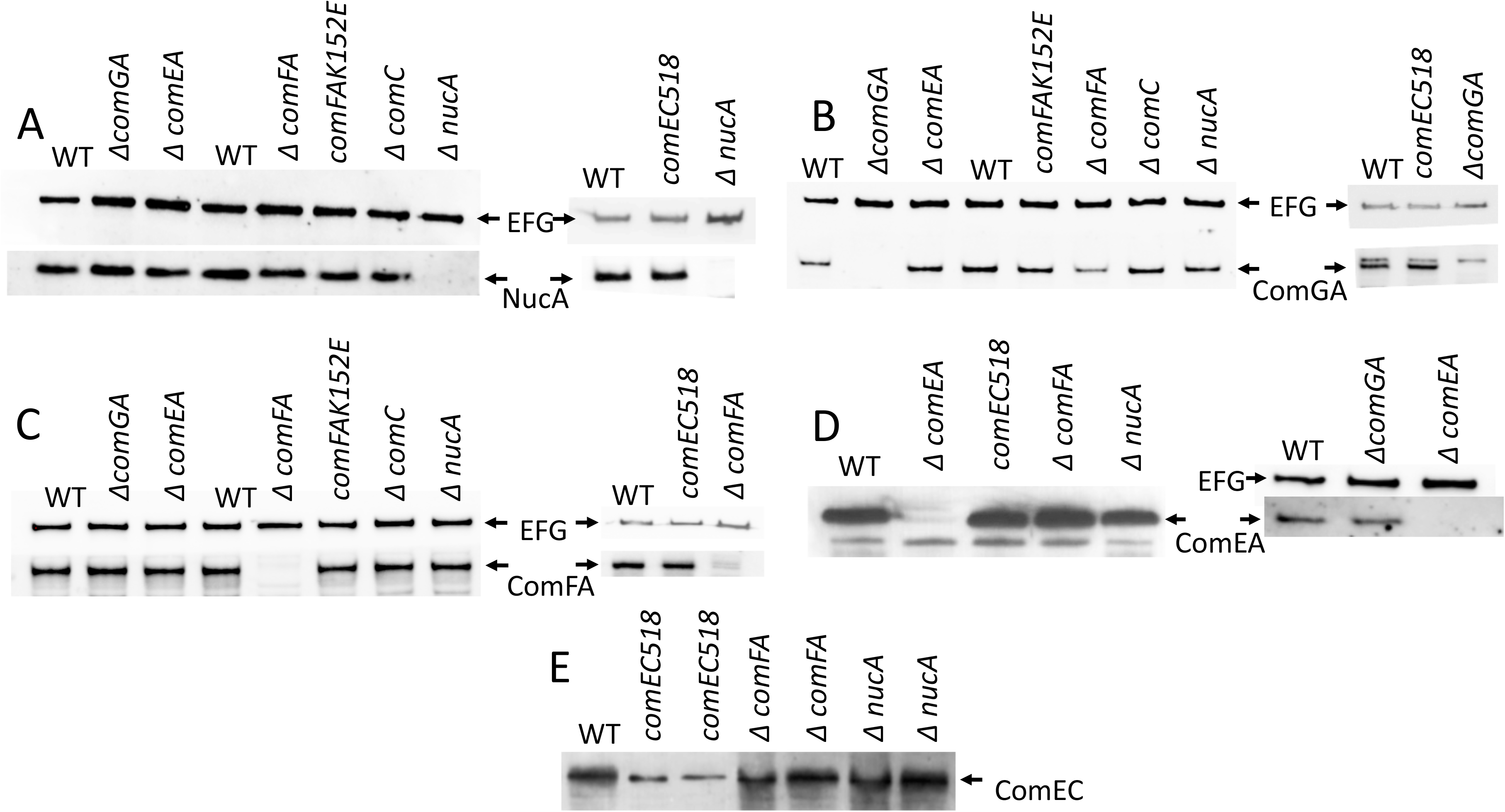
Western blots for transformation proteins in mutant strains. All of the mutants were deletions except for *comEC518* which is a transposon insertion near the beginning of the coding sequence. In all cases the relevant knockout mutant was used as a negative control for the antiserum. There is a consistent cross-reacting band at the position of the ComEC signal. For most of the gels a loading control is shown using anti-elongation factor G (EFG) antiserum, a kind gift from J. Dworkin. In panel D, the cross-reacting band just below the ComEA signal serves as a loading control. In panel E, a cross-reacting band is visible in the *comEC518* lanes at the position of ComEC (29).

**Figure S2.**
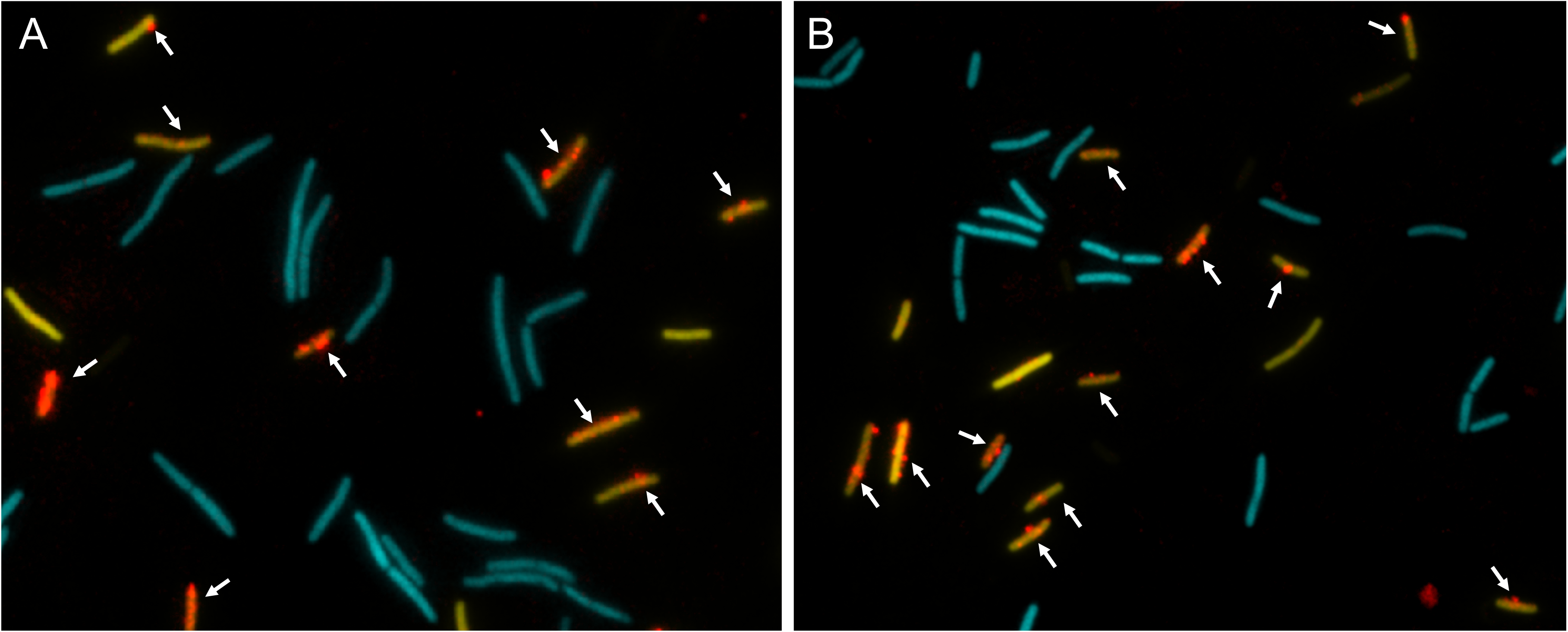
DNA binding in Δ*comGA* and Δ*comC* strains. Δ*comGA* (A) and Δ*comC* (B) strains, expressing CFP, were combined with transformed wild-type bacteria expressing YFP. Transformation was for 30 minutes without DNAase treatment. The arrows indicate all the cells with detectable rDNA signals. The CFP labeled Δ*comGA* cells are slightly filamented (Hahn J, Tanner AW, Carabetta VJ, Cristea IM, Dubnau D. 2015. Mol. Microbiol 97:454-71).

**Figure S3.**
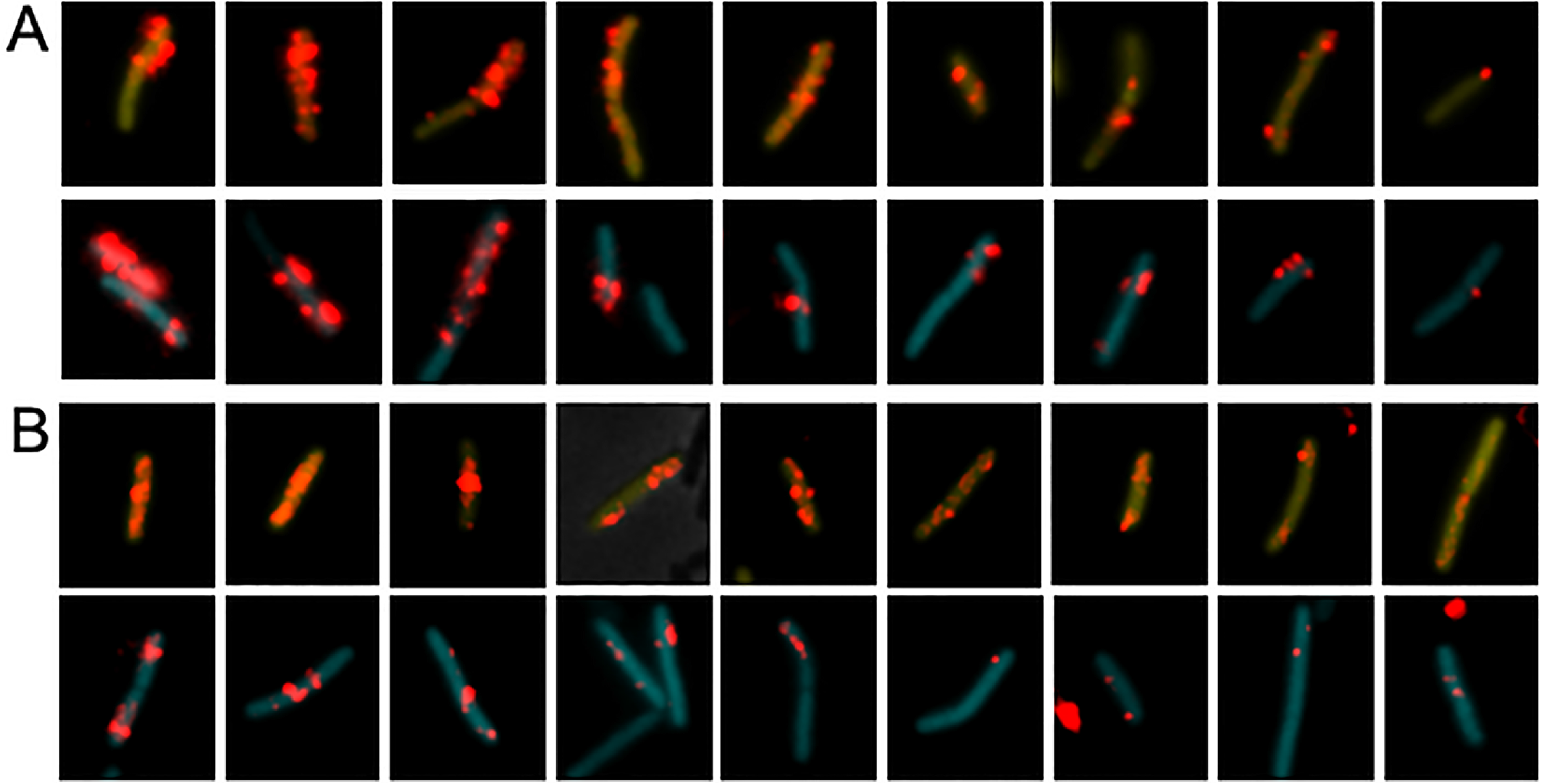
Binding and uptake in the Δ*comFA* mutant. Wild-type (YFP) and mutant (CFP) cells were combined before 30-minutes incubation with rDNA. Panels A and B show results for Δ*comFA* without and with DNAase treatment respectively. In each panel the top and bottom rows show 9 each of wild-type and mutant cells respectively. All the cells were taken from a single representative microscope field and show all or nearly all of the rDNA-associated cells in that field. In the cropped images of a given panel, rDNA signals were enhanced identically but rDNA signal intensities cannot be compared between panels A and B.

**Figure S4.**
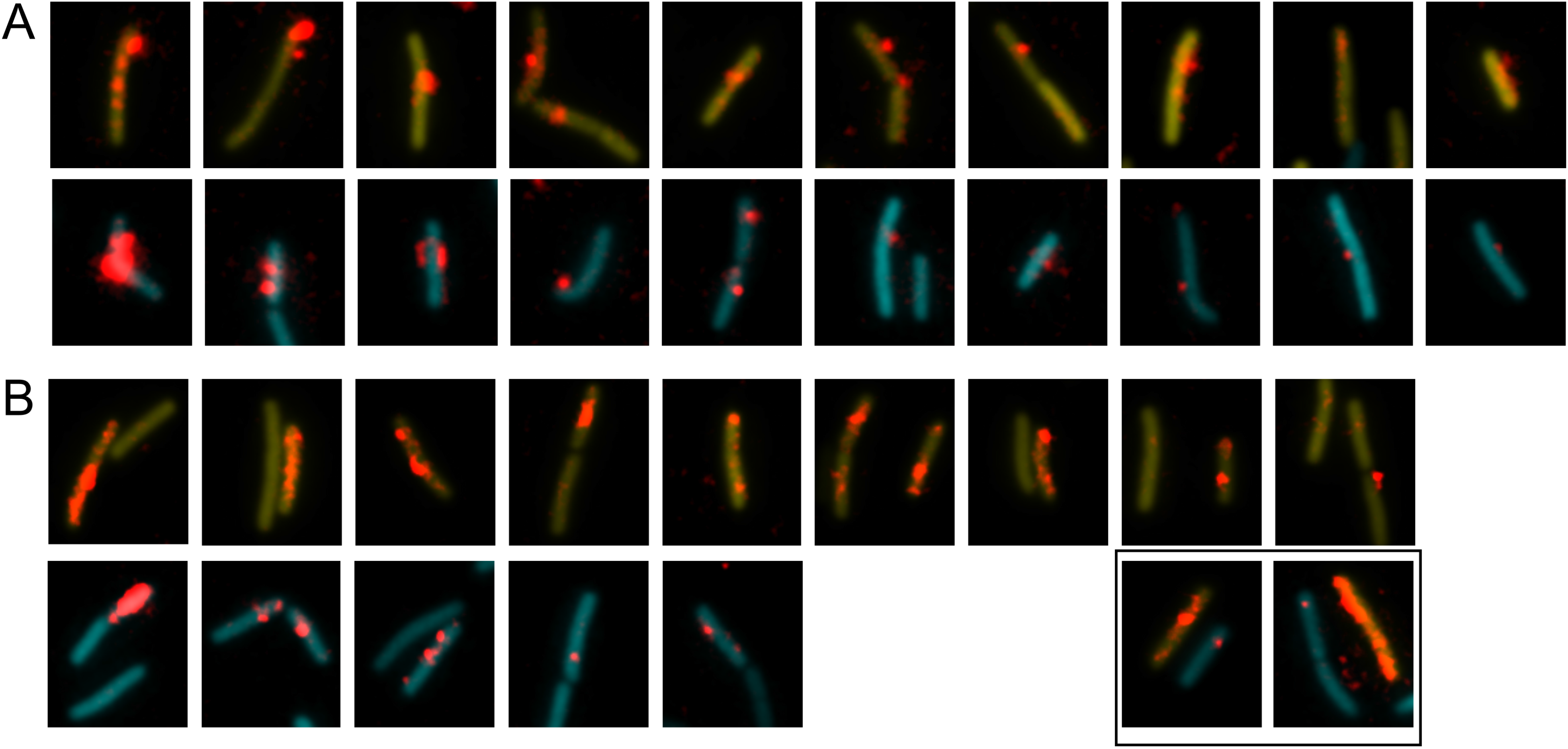
Binding and uptake in the Δ*nucA* mutant. Wild-type (YFP) and mutant (CFP) cells were combined before 15-minutes incubation with rDNA and imaged without (panel A) and with DNAase (panel B). In each of these panels the top and bottom rows show nearly all the wild-type and mutant cells from a single field, respectively. The rhodamine signal was enhanced before cropping the cells and the intensities of the rhodamine signal can be directly compared. The YFP and CFP images were separately adjusted in each image so as not to obscure the rDNA signal. The images in these two panels were ordered by apparent rDNA signal strengths decreasing from left to right, to facilitate comparisons of the mutant and wild-type cells. The boxed images both a YFP- and a CFP-expressing cell.

**Figure S5.**
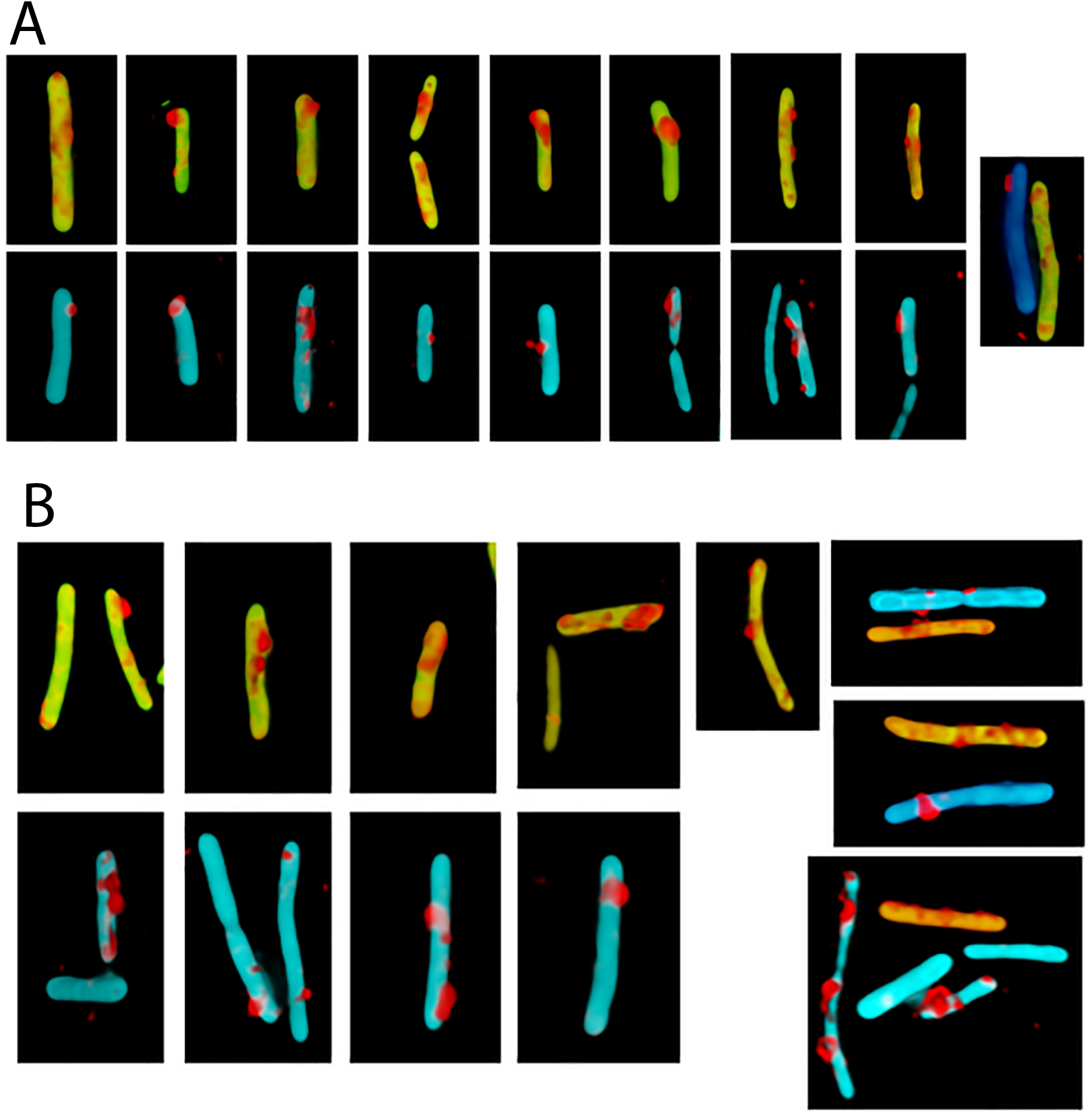
Volume reconstructions of the Δ*nucA* (panel A) and *nucA* D98A (panel B) mutant strains. Wild-type (YFP-expressing) and mutant (CFP-expressing) cells are shown in each panel for comparison.

