## Supplemental material for "Mechanisms of transforming DNA uptake to the periplasm of *Bacillus subtilis*"

Table S1

Strains

| **Strain**[a] | **Genotype** | **Source** |
|---|---|---|
| BD5810 | *amyE::*$P_G$*-CFP (spc)* | (48) |
| BD6011 | *amyE::*$P_G$*-YFP (spc)* | (48) |
| BD8739 | t*rpC2 ΔcomEA (ery)* [b] | |
| BD8800 | *amyE::Pr*$_G$*-CFP (spc) ΔcomGA(ery)*[b] | This work |
| BD8802 | *amyE:: Pr*$_G$*-CFP (spc) ΔcomEA(ery)* [b] | This work |
| BD8805 | *amyE::*$P_G$*-CFP (spc) ΔnucA(ery)* [b] | This work |
| BD8883 | *amyE::*$P_G$*-CFP (spc) ΔcomC (ery)* [b] | This work |
| BD8884 | *amyE::*$P_G$*-CFP (spc) comFA K152E* | (33) |
| BD8911 | *amyE::*$P_G$*-CFP (spc) ΔcomFA (ery)* [b] | This work |
| BD8949 | *amyE::*$P_G$*-CFP (spc) yfp-comEA ΔcomEA (ery)* [b] | This work |
| BD8951 | *amyE::*$P_G$*-CFP (spc) nucAD98A* | This work |
| BD8954 | *amyE::*$P_G$*-CFP (spc) comEC-518* | (29) |

[a]All of the listed strains except BD8739 were derivatives of IS75 *his leu met*.

[b]These deletion mutations, constructed by Koo et al (63), were obtained from the Bacillus Genetic Stock Center

Table S2

Primers

| Primer number | Sequence (5’->3’) |
|---|---|
| 1 | AGAGGATCCCCGGGTACCGAGCTCGAATTCAAAAGGTATCGAGTAGCCAG |
| 2 | CTCCTTTACTCATATTCATCGTGCATGTTCC |
| 3 | ATGCACGATGAATATGAGTAAAGGAGAAGAAC |
| 4 | TCAACCAATTCATTTTGTATAGTTCATCCATGC |
| 5 | GGAAACAGCTATGACCATGATTACGAATTCACTTTACTGTAATGGAAG |
| 6 | ACTATACAAATAAATGAATTGGTTGAATCAGC |

Figure S1. Western blots for transformation proteins in mutant strains. All of the mutants were deletions except for *comEC518* which is a transposon insertion near the beginning of the coding sequence. In all cases the relevant knockout mutant was used as a negative control for the antiserum. There is a consistent cross-reacting band at the position of the ComEC signal. For most of the gels a loading control is shown using anti-elongation factor G (EFG) antiserum, a kind gift from J. Dworkin. In panel D, the cross-reacting band just below the ComEA signal serves as a loading control. In panel E, a cross-reacting band is visible in the *comEC518* lanes at the position of ComEC (29).

Figure S2. DNA binding in *ΔcomGA* and *ΔcomC* strains. *ΔcomGA* (A) and *ΔcomC* (B) strains, expressing CFP, were combined with transformed wild-type bacteria expressing YFP. Transformation was for 30 minutes without DNAase treatment. The arrows indicate all the cells with detectable rDNA signals. The CFP labeled *ΔcomGA* cells are slightly filamented (Hahn J, Tanner AW, Carabetta VJ, Cristea IM, Dubnau D. 2015. Mol. Microbiol 97:454-71).

Figure S3. Binding and uptake in the *ΔcomFA* mutant. Wild-type (YFP) and mutant (CFP) cells were combined before 30-minutes incubation with rDNA. Panels A and B show results for *ΔcomFA* without and with DNAase treatment respectively. In each panel the top and bottom rows show 9 each of wild-type and mutant cells respectively. All the cells were taken from a single representative microscope field and show all or nearly all of the rDNA-associated cells in that field. In the cropped images of a given panel, rDNA signals were enhanced identically but rDNA signal intensities cannot be compared between panels A and B.

Figure S4. Binding and uptake in the *ΔnucA* mutant. Wild-type (YFP) and mutant (CFP) cells were combined before 15-minutes incubation with rDNA and imaged without (panel A) and with DNAase (panel B). In each of these panels the top and bottom rows show nearly all the wild-type and mutant cells from a single field, respectively. The rhodamine signal was enhanced before cropping the cells and the intensities of the rhodamine signal can be directly compared. The YFP and CFP images were separately adjusted in each image so as not to obscure the rDNA signal. The images in these two panels were ordered by apparent rDNA signal strengths decreasing from left to right, to facilitate comparisons of the mutant and wild-type cells. The boxed images both a YFP- and a CFP-expressing cell.

Figure S5. Volume reconstructions of the *ΔnucA* (panel A) and *nucA* D98A (panel B) mutant strains. Wild-type (YFP-expressing) and mutant (CFP-expressing) cells are shown in each panel for comparison.

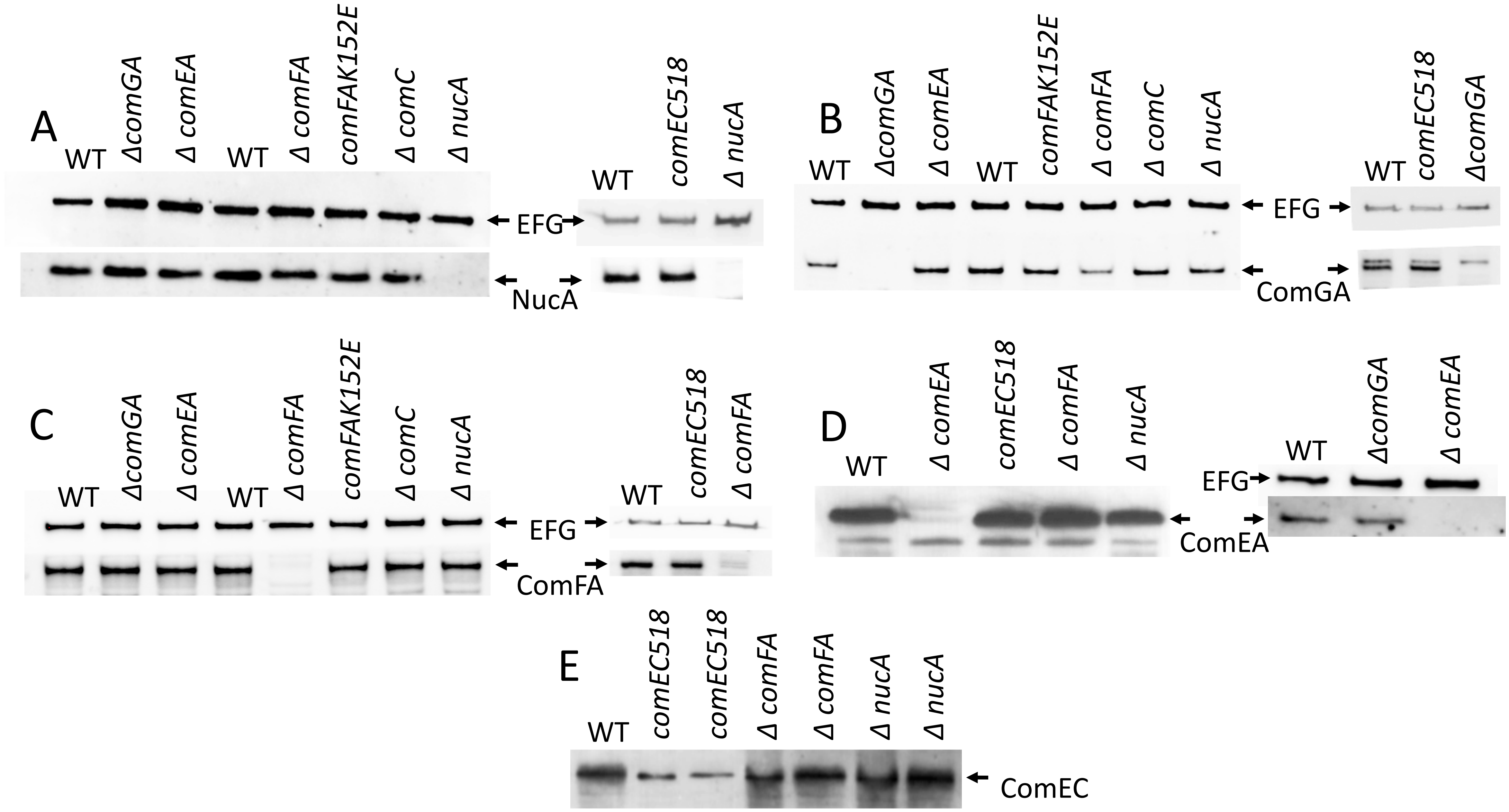
A
WT
ΔcomGA
Δ comEA
WT
Δ comFA
comFAK152E
Δ comC
Δ nucA
EFG
NucA
WT
comEC518
Δ nucA
B
WT
ΔcomGA
Δ comEA
WT
comFAK152E
Δ comFA
Δ comC
Δ nucA
EFG
ComGA
WT
comEC518
ΔcomGA
C
WT
ΔcomGA
Δ comEA
WT
Δ comFA
comFAK152E
Δ comC
Δ nucA
EFG
ComFA
WT
comEC518
Δ comFA
D
WT
Δ comEA
comEC518
Δ comFA
Δ nucA
ComEA
WT
ΔcomGA
Δ comEA
EFG
E
WT
comEC518
comEC518
Δ comFA
Δ comFA
Δ nucA
Δ nucA
ComEC

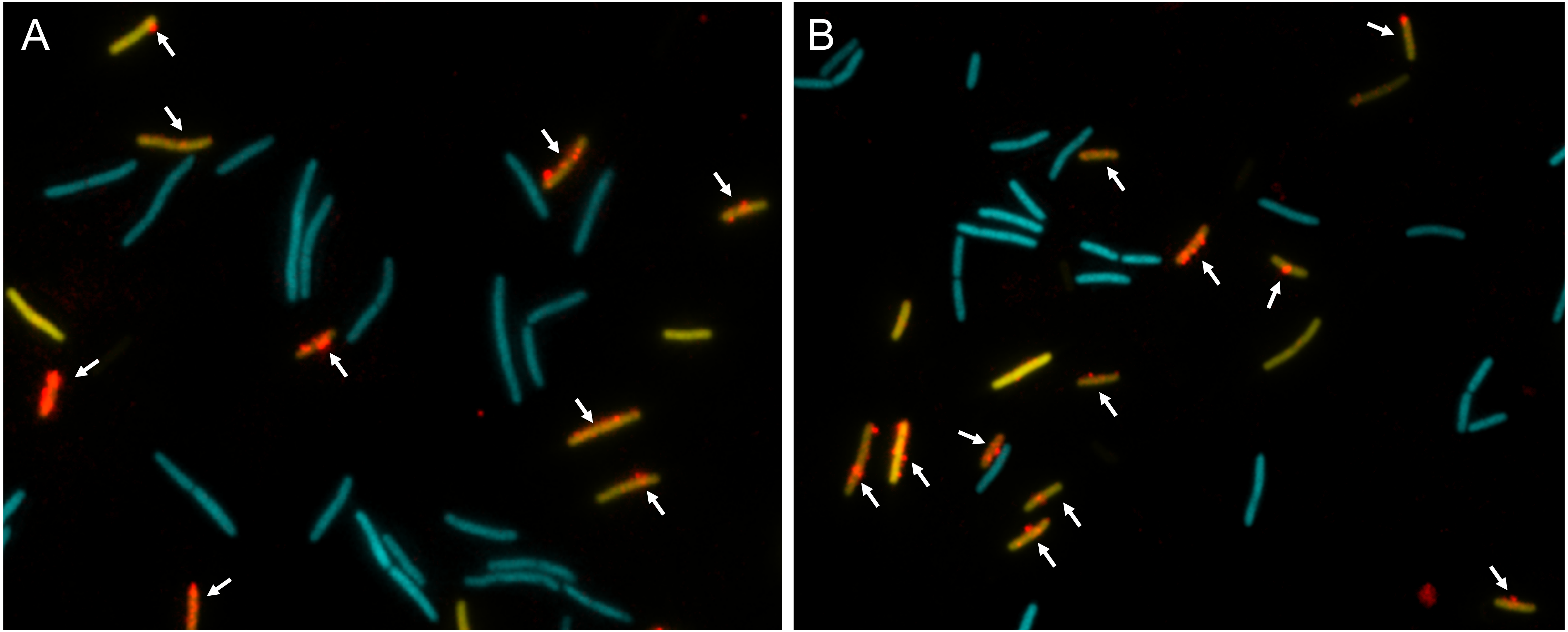
A
B

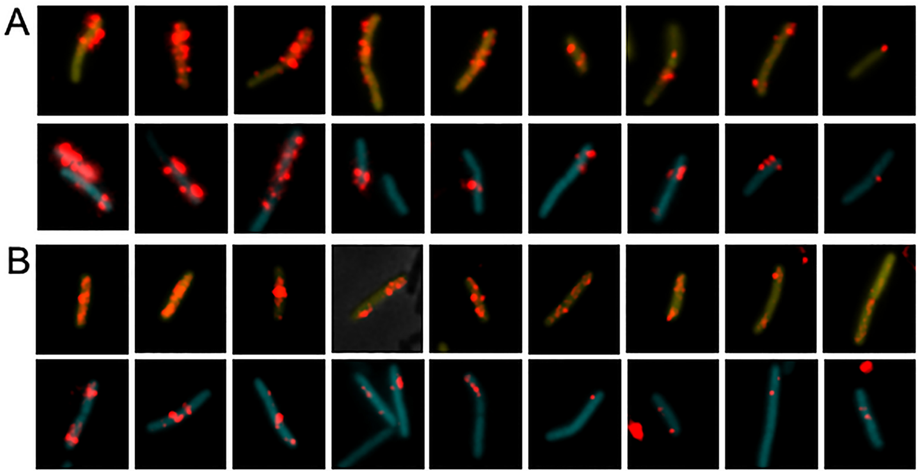
A
B

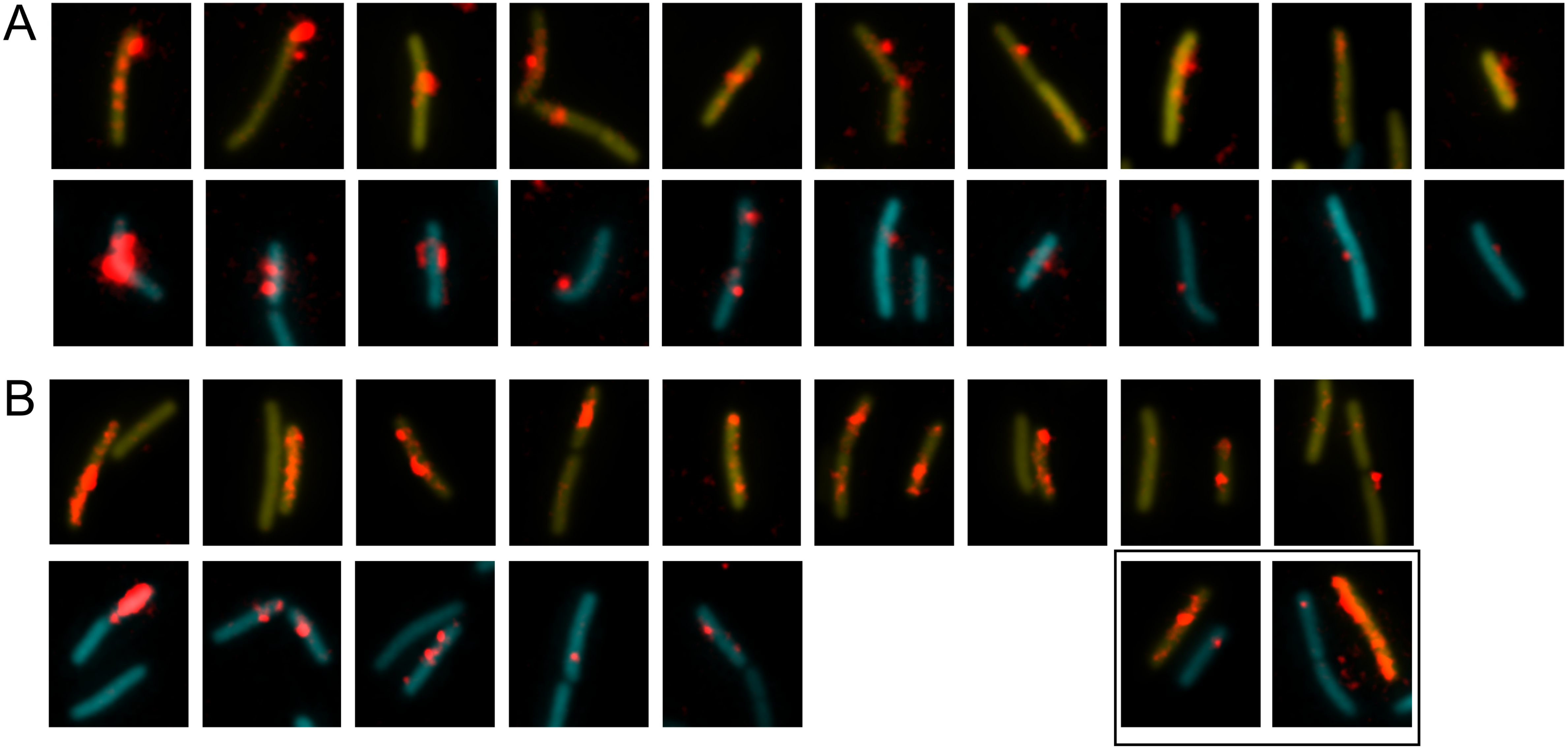
A
B

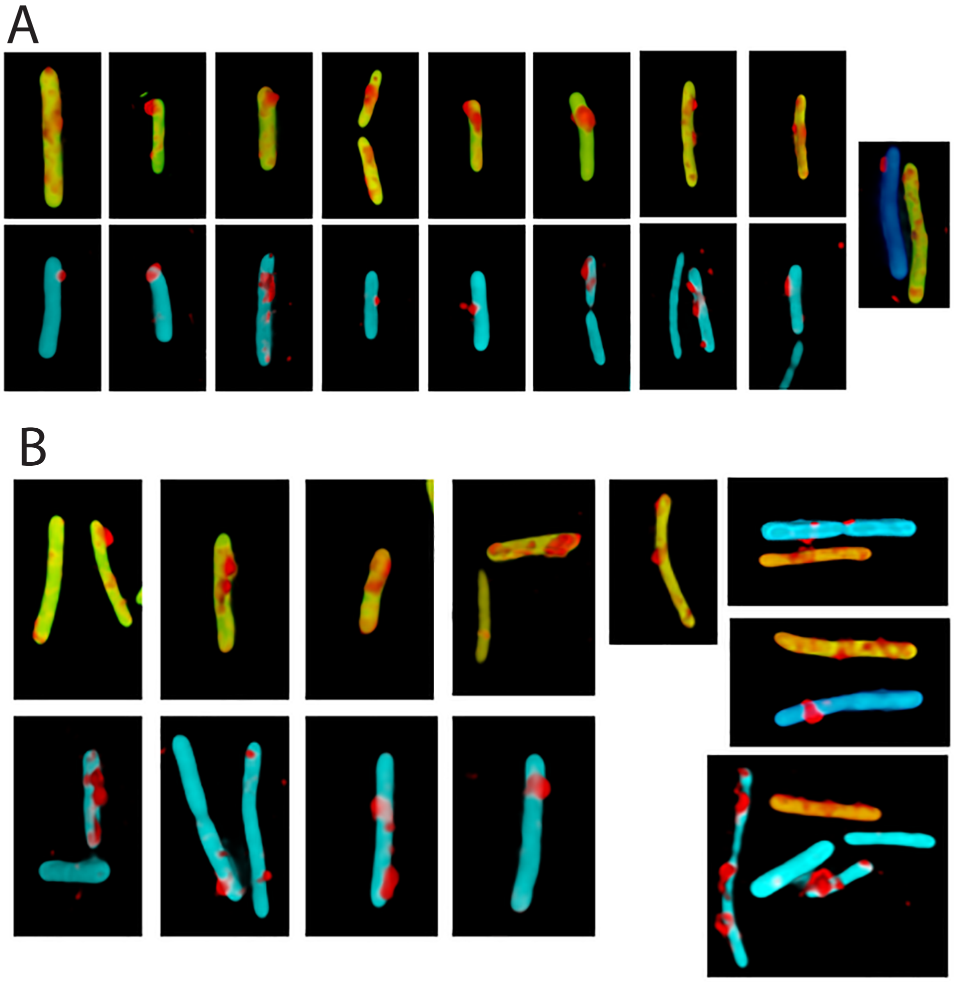
A
B
